# Clearance of genome-damaged cells from the hematopoietic system via p53 without contribution by the cGAS/STING axis

**DOI:** 10.1101/2022.06.24.497496

**Authors:** Nicole Dressel, Loreen Natusch, Clara M. Munz, Santiago Costas Ramon, Mina N.F. Morcos, Anja Loff, Björn Hiller, Mathias Lesche, Andreas Dahl, Hella Luksch, Angela Rösen-Wolff, Axel Roers, Rayk Behrendt, Alexander Gerbaulet

**Affiliations:** Institute for Immunology, Faculty of Medicine, TU Dresden, 01307 Dresden, Germany; DRESDEN-concept Genome Center, Center for Molecular and Cellular Bioengineering, TU Dresden, 01307 Dresden, Germany; Department of Pediatrics, University Hospital Carl Gustav Carus, TU Dresden, 01307 Dresden, Germany; Institute for Immunology, Heidelberg University Hospital, 69120 Heidelberg, Germany; Institute for Clinical Chemistry and Clinical Pharmacology, University Hospital Bonn, 53127 Bonn, Germany

## Abstract

Cell-intrinsic response patterns control risks arising from genome-damage, preventing malignant transformation. The DNA sensor cyclic-GMP-AMP synthase (cGAS) has emerged as a new principle detecting genome damage, as it can be triggered by aberrant self-DNA. Stimulator of interferon genes (STING)-activation downstream of cGAS can drive cells into senescence or cell death and induces antiproliferative type I interferon (IFN) and pro-apoptotic tumor necrosis factor responses. Herein, we investigated how DNA damage-driven activation of cGAS/STING signaling impacts on hematopoiesis. Defective ribonucleotide excision repair (RER) in the hematopoietic system caused chromosomal instability as well as robust activation of the cGAS/STING/IFN axis, and compromised hematopoietic stem cell function, resulting in cytopenia and ultimately leukemia. Whereas loss of p53 largely rescued RER-deficient hematopoiesis at the cost of further accelerated leukemogenesis, the additional inactivation of cGAS, STING or type I IFN signaling had no detectable effect on blood cell generation and leukemia development. Moreover, cGAS-deficient hematopoiesis showed unaltered responses to spontaneous or acute DNA damage. Our data demonstrate that the cGAS/STING pathway is dispensable for the hematopoietic system coping with chronic or acute DNA damage and does not protect against leukemic transformation in the absence of RER.

## Introduction

Genome integrity is continuously challenged by DNA damage resulting from spontaneous hydrolysis of phosphodiester bonds, ionizing radiation, mutagenic chemicals or reactive oxygen species (Lindahl, 2016). Cells detect DNA damage and activate signaling cascades to halt the cell cycle and induce repair pathways (Jackson and Bartek, 2009). The tumor suppressor p53 is key in cellular responses to diverse forms of stress, including genome damage. DNA damage signaling stabilizes the p53 protein, allowing it to regulate transcription of numerous genes. Depending on quality of DNA damage, signal intensity and cell type, p53 mediates cell cycle arrest, triggers DNA repair, activates different forms of programmed cell death or drives cells into senescence (Hafner et al., 2019; Ou and Schumacher, 2018; Vousden and Prives, 2009). Persistent DNA damage that cannot be repaired has fundamentally different consequences on short lived, post-mitotic cells, in contrast to long-lived stem and progenitor cells with high proliferative potential (Mandal et al., 2011).

Genome damage was recently demonstrated to activate cell-intrinsic type I interferon (IFN) production (Erdal et al., 2017; Hartlova et al., 2015; Hiller et al., 2018; Li and Chen, 2018; Mackenzie et al., 2017). An important link between genome damage and innate antiviral immunity is the pattern recognition receptor cyclic GMP-AMP synthase (cGAS), which detects double-stranded DNA. Upon ligation by DNA, cGAS catalyzes formation of a second messenger, cyclic guanosine monophosphate–adenosine monophosphate (cGAMP) that in turn activates the sensor stimulator of interferon genes (STING) to induce type I IFN and proinflammatory cytokine production. In addition to viral DNA, cGAS can be activated by endogenous self-DNA (Hopfner and Hornung, 2020), including cytosolic chromatin resulting from genotoxic stress (Dou et al., 2017; Glück et al., 2017; Harding et al., 2017; Mackenzie et al., 2017) and release of mitochondrial DNA (West et al., 2015), thereby triggering type I IFN responses (Ahn et al., 2014; Erdal et al., 2017; Schubert et al., 2022; Shen et al., 2015; Takahashi et al., 2018; Vanpouille-Box et al., 2017). Type I IFN exerts robust antiproliferative effects on several types of more mature cells (Bromberg et al., 1996; McNab et al., 2015; Sangfelt et al., 2000). In addition, cGAS/STING signaling can also trigger senescence and cell death (Glück et al., 2017; Gulen et al., 2017; Li and Chen, 2018; Paludan et al., 2019), potentially eliminating genome-damaged cells and thereby preventing cancer (Li and Chen, 2018).

Herein, we address the effects of DNA damage-induced cGAS/STING responses *in vivo*. We use mice with hematopoieitic deficiency of the enzyme RNase H2 (RH2) as a model for chronic DNA damage. RH2 is essential for the ribonucleotide excision repair (RER) pathway responsible for removal of vast numbers of single ribonucleotides that the replicative polymerases misincorporate into genomic DNA during S-phase of the cell cycle (Hiller et al., 2012; Reijns et al., 2012b). Failure to repair these lesions results in genome instability, micronucleus formation, chronic activation of cGAS/STING and type I IFN production (Bartsch et al., 2017; Hiller et al., 2018; Mackenzie et al., 2016; Mackenzie et al., 2017; Reijns et al., 2012a). We find that the genome damage ensuing from loss of RER severely compromised hematopoiesis and resulted in malignant transformation. Additional loss of p53 largely rescued blood cell production at the cost of further accelerated leukemogenesis. Unexpectedly, inactivation of the cGAS/STING axis or of type I IFN signaling had no detectable impact on hematopoiesis and leukemia development in RER-deficient mice. In addition, exclusive loss of cGAS/STING did neither alter steady state nor stress hematopoiesis. Our findings argue that the cGAS/STING axis does not significantly contribute to protection of the hematopoietic system against DNA damage accumulation and malignant transformation.

## Methods

Detailed information about cell preparation, flow cytometry, bone marrow transplantation, HSC culture, histology, gene expression analysis and statistics are provided in the Supplemental methods.

### Mice

The following mouse strain were used in the study: *Rnaseh2b*^flox^ (Hiller et al., 2012), Vav-Cre (Stadtfeld and Graf, 2005), *Ifnar1*^KO^ (Kamphuis et al., 2006), *Cgas*^KO^ (Schoggins et al., 2014), *Trp53*^KO^ (Donehower et al., 1992), *Sting1*^GT^ (Sauer et al., 2011) and B6.CD45.1 (RRID:IMSR_JAX:002014). Male and female mice were used for experiments and housed in individually ventilated cages under specific-pathogen free environment at the Experimental Center of the Medical Faculty, TU Dresden. 5-fluorouracil (150 µg/g body weight, Applichem) was administered via intravenous (i.v.) injection, whole body γ-radiation was applied using a MaxiShot source (Xylon). All animal experiments were in accordance with institutional guidelines and were approved by the Landesdirektion Dresden.

### PB erythrocyte micronucleus assay

Micronucleated erythrocytes were identified following the protocol of Balmus et al. (Balmus et al., 2015). Briefly, 30µl PB was mixed with 120µl PBS and fixed with pre-cooled (−80°C) methanol overnight and stored for <1 year at -80%. Fixed erythrocytes were stained with antibodies against CD71 and Ter119 and digested with 10µl RNase A (10mg/ml, Invitrogen) for 1hour at 4°C. Samples were washed and after adding 1µg/ml propidium iodide (PI), 1-2.5 × 10^6^ events were acquired on BD LSR II or BD FACS Aria III flow cytometers. Representative gating of micronucleated erythrocytes is shown in Figure S1E.

### Statistics

Statistical analysis was performed using Graphpad Prism 9 and applied tests are given in each figure legend (ns= not significant, * = p <0.05 - 0.01, ** = p <0.01 – 0.001, *** = p < 0.001).

## Results

### Hematopoietic loss of RER results in genome instability and predisposes to leukemia

In order to investigate defense-mechanisms against genome damage in the hematopoietic system, we generated mice with loss of RER throughout hematopoiesis by conditional inactivation of the *Rnaseh2b* gene (Hiller et al., 2012; Stadtfeld and Graf, 2005). As expected, *Rnaseh2b*^FL/FL^/Vav-Cre^+^ mice, (“RH2^hKO^”) featured high numbers of micronucleated erythrocytes and histone 2AX-phosphorylated (γH2AX) bone marrow (BM) cells (Figures 1A-D, S1) reflecting genome instability ensuing from unrepaired ribonucleotides contained in the genomic DNA (Balmus et al., 2015; Cerritelli and Crouch, 2016; Kellner and Luke, 2020; Williams et al., 2016). Except for a minor growth retardation (Figure S2A), the mutants are macroscopically indistinguishable from Cre-negative control littermates (“RH2^hWT^”) during the first 3 months of life. However, median survival was significantly reduced to about 13 months (Figure 1E). Analysis of moribund mice at different age revealed a massive enlargement of the thymus (Figures 1F-G). Histology showed a loss of normal thymic medulla and cortex architecture (Figure S2B) due to a massive expansion of CD4/8 double positive (DP) thymocytes (Figure 1H). In some of these animals, DP thymocytes disseminated to PB (Figure 1I) and spleen (Figure S2B), suggesting T cell acute lymphoblastic leukemia (T-ALL)-like disease as the most likely cause of death in these animals. Some moribund mice did not show signs of leukemia as judged by macroscopic inspection of organs or immuno-phenotyping of PB, BM, thymus and spleen. These animals presented with very low hematocrit and leukocytes counts, suggesting hematopoietic failure or infection as the reason for lethality.

**Figure 1.**
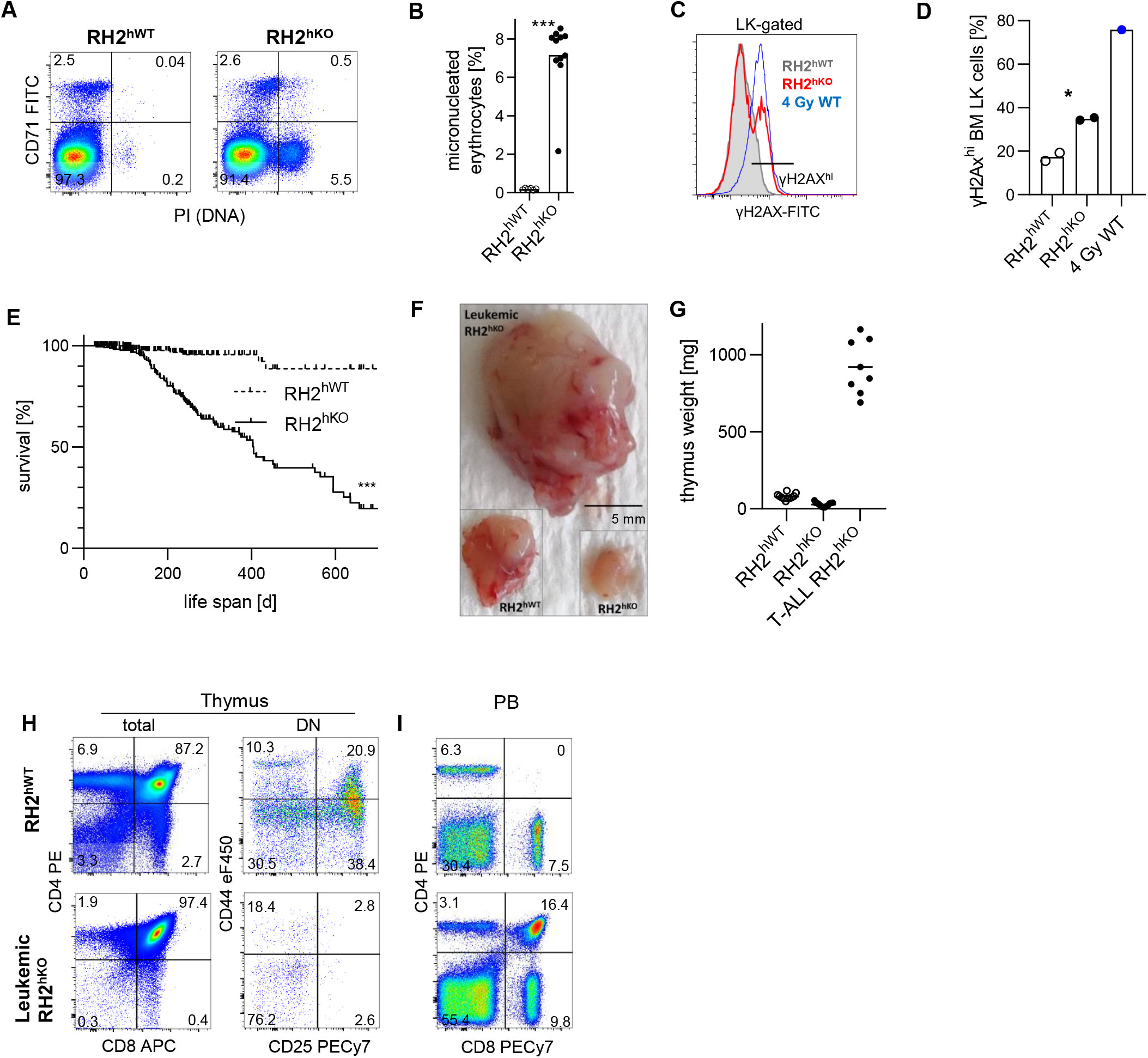
Hematopoietic loss RER results in genome instability and predisposes to leukemia. **A** Detection of micronucleated erythrocytes in peripheral blood (PB) of animals with hematopoietic RNase H2 deficiency (RH2^hKO^, right) or Cre-negative controls (RH2^hWT^, left); representative examples of the data in B. **B** Frequency of micronucleated erythrocytes among normochromic erythrocytes (n=6-11/genotype, individual mice (15-47 wks of age) and means (bars) are shown, significance was calculated by an unpaired Student’s t test). **C** Intranuclear detection of γH2AX-phosphorylation in lineage^-^ CD117^hi^ (LK) BM cells isolated from RH2^hWT^ and RH2^hKO^ mice (n=2/genotype, representative example of data in D). A WT mouse was exposed to 4 Gy γ-radiation 90 min before analysis and served as a positive control. **D** Percentage of γH2AX^hi^ LK BM cells (unpaired Student’s t test, individual mice and means shown (bars)). **E** Kaplan-Meier survival curve of RH2^hKO^ animals and RH2^hWT^ littermate controls. Significance was calculated by log-rank test. **F** Representative example of an enlarged thymus from a leukemic RH2^hKO^ mouse; insets show thymi from either RH2^hWT^ control or non-leukemic RH2^hKO^ animals at the same magnification. **G** Thymus weight of RH2^hWT^ (n=11), non-leukemic RH2^hKO^ (n=13) and leukemic (T-ALL) RH2^hKO^ (n=8) mice (aged 6-36 wks). **H-I** Representative flow cytometric analysis of total (left) and CD4/8 double-negative (DN, right) thymocytes (H) and peripheral blood (PB, I) isolated from a leukemic RH2^hKO^ (lower row) and an RH2^hWT^ control (upper row) mouse.

These data show that conditional inactivation of RER represents a suitable model for studying hematopoiesis under conditions of chronic DNA damage.

### High load of genome damage causes hematopoietic malfunction in RH2^hKO^ mice

At young age before onset of malignancy, RH2^hKO^ animals featured massive alterations of the hematopoietic system. These included macrocytic anemia, almost complete absence of B cells, reduced T cell and granulocyte numbers, but elevated platelet counts (Figure 2A). Analysis of RH2^hKO^ BM revealed that total cellularity was reduced to about one third of controls. To robustly identify hematopoietic stem cells (HSCs, see Figure S1A for gating) we substituted the marker Sca-1, which is upregulated by type I IFN signaling (Kanayama et al., 2020) with EPCR (CD201) (Figure 2B)(Vazquez et al., 2015). Like HSCs, also absolute numbers of progenitor populations were massively reduced in RH2^hKO^ BM (Figure 2B). All populations of B-lymphocyte development from pre-pro B cells to mature B cells (see Figure S1B for gating) were reduced to below 10% of normal numbers (Figure 2C). Likewise, the thymi of RH2^hKO^ mice were severely reduced in weight (Figures S3A-B) and cellularity was only about 5% of controls (Figure 2D). Thymocytes of the first stages of T lymphocyte development (CD4/8 double-negative (DN) 1-3), see Figure S1C for gating) were reduced to about half of normal numbers and cells of later stages were almost completely absent (Figure 2D). About 10% of the RH2^hKO^ mice distinctly differed from the majority by ameliorated cytopenia (Figures S3C-F) and showed almost normal numbers of micronucleated erythrocytes (Figure S3G), but retained high Sca-1 expression (Figure S3H), most likely reflecting overgrowth of clones that had escaped Cre-mediated inactivation of *Rnaseh2b*.

**Figure 2.**
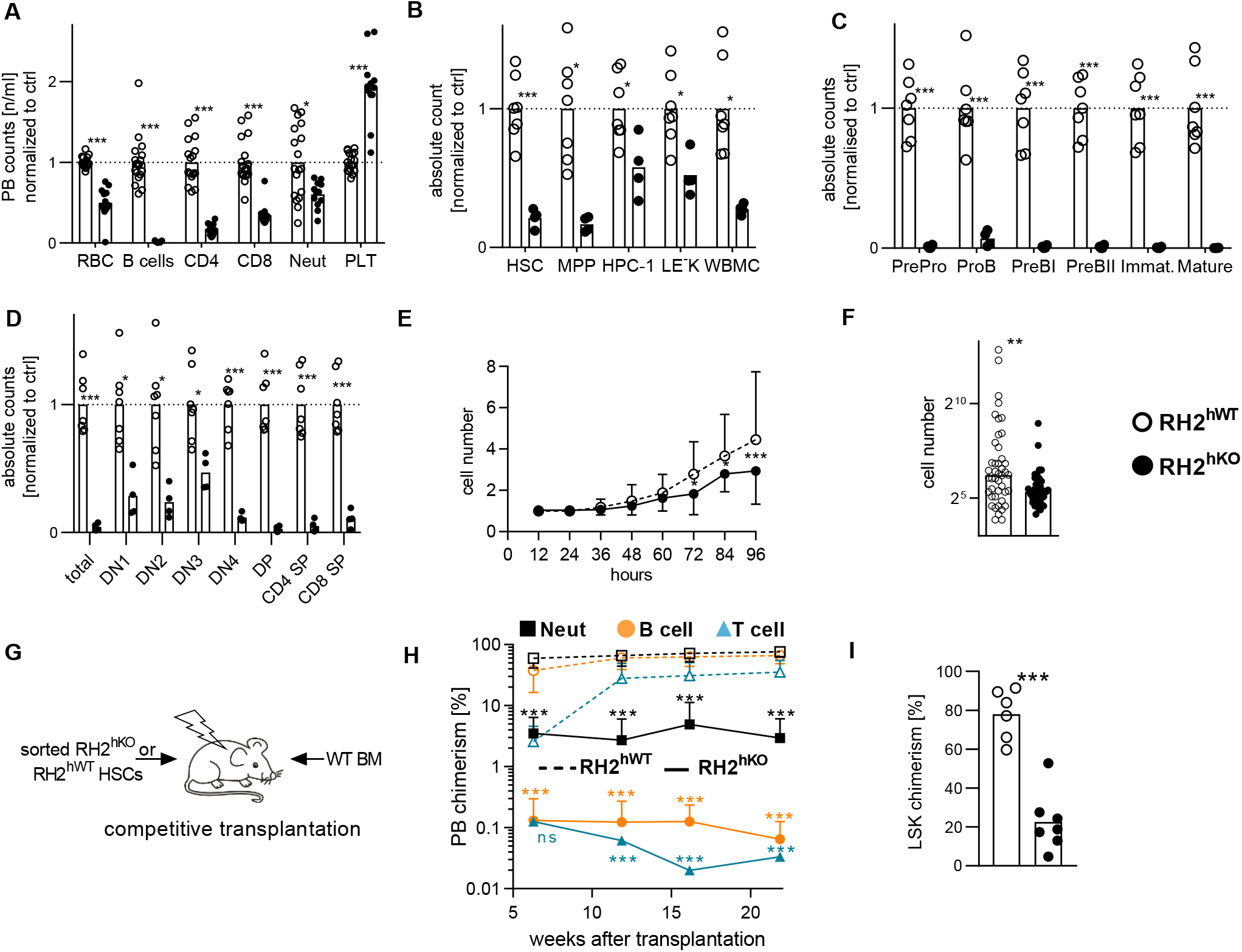
Chronic genome damage impairs blood cell production and HSC fitness. **A** PB cell counts of RH2^hKO^ (closed circles) and Cre-negative controls (RH2^hWT^, open circles). Erythrocyte (RBC), B cell, CD4/CD8 T lymphocyte, neutrophil (Neut), and platelet (PLT) numbers were determined (n=13-18/genotype, individual mice and means (bars) are shown, cell numbers were normalized to RH2^hWT^ ctrl means (set to 1, dotted line), unpaired Student’s t test with Holm-Sidak post test). **B** Absolute numbers of bone marrow hematopoietic stem and progenitor cells (BM HSPCs) were determined (HSCs, LEK CD48^-/lo^CD150^+^), multipotent (MPP, LEK CD48^-^CD150^-^) hematopoietic progenitor cells HPC-1 (LEK CD48^+^CD150^-^), erythro-myeloid progenitors (LE^-^K) progenitors and whole bone marrow cellularity (WBMC) were determined (n=4-7/genotype, display, normalization and statistic analysis of data as in A). **C** Absolute numbers of developing B lymphocyte progenitors in the bone marrow. (n=4-7/genotype; display, normalization and statistic analysis of data as in A). **D** Thymocyte development (DN, CD4/CD8 double-negative; DP, double-positive; SP, single-positive; n=4-7/genotype; display, normalization and statistical analysis of data as in A). **E-F** Single HSCs (LSK CD48^-/lo^CD150^+^CD34^-/lo^CD201^hi^, n=38-44) isolated from either RH2^hKO^ or RH2^hWT^ mice were cultivated *in vitro*. (E) Cells were counted for 96h every 12h, mean & SD are shown, significance was calculated by repeated measures 2-way ANOVA with Sidak post test). (F) Colony size after 10 days of culture (median and individual colonies are shown, significance was calculated by Mann-Whitney U test).**G-I** 200 RH2^hKO^ (closed symbols, solid line) or RH2^hWT^ (open symbols, dotted line) HSCs (LEK CD48^-/lo^CD150^+^) were mixed with 500,000 WT B6.CD45.1 WBMCs and competitively transplanted into lethally irradiated B6.CD45.1/CD45.2 recipient mice (G, n=6-7/genotype). Donor chimerism of PB neutrophils (Neut, square), B (circle) and T (triangle) cells (H, mean and SD is shown, significance was calculated by repeated measures 2-way ANOVA with Sidak post test) and BM LSK cells (I, individual mice and means (bars) are shown, significance was calculated by an unpaired Student’s t test) was quantified.

To address stem cell fitness, we cultivated single HSCs and found a reduced proliferative capacity of RH2^hKO^ HSCs (Figures 2E-F). Moreover, we determined the functional potential of HSCs *in vivo* by competitive transplantation into lethally irradiated recipients and found a severe repopulation defect of RNaseH2-deficient HSCs (Figures 2G-I, S2I-J).

Taken together, chronic genome damage in the hematopoietic system of RH2^hKO^ mice reduced numbers, proliferation and repopulation capacity of HSCs. Impaired hematopoiesis resulted in anemia, lymphopenia and reduced granulocyte numbers.

### Genome damage in RH2^hKO^ mice activates p53 and type I IFN signaling

Inflammatory conditions confound the fidelity of HSPC marker expression (Kanayama et al., 2020), but granulocyte-monocyte progenitors (GMPs) are unambiguously identified by upregulation of Fcγ receptors (CD16/32). Moreover, genome-damage related to loss of RER primarily manifests in replicating cells (Williams et al., 2016) and GMPs are rapidly dividing cells. In order to elucidate how chronic genome damage leads to hematopoietic malfunction in RH2^hKO^ mice, we sorted GMPs from BM of mutant and control animals and sequenced their transcriptomes. We found 198 genes significantly up-regulated in RH2^hKO^ GMPs compared to controls (Figure 3A, Table S1). Among the most upregulated genes were the ISGs *Oasl1, Mx1, Ifit1* (Figure 3B, Table S2) as well as *Cdkn1a*, encoding cell cycle inhibitor p21 and other p53-induced genes, including *Trp53inp1, Bax* and *Mdm2 (*Figure 3C, Table S3*). Rnaseh2b* was the most downregulated gene, validating the analysis. In addition, *Ear1/2, Gzmb* and *Tspan9* were among 20 significantly downregulated genes. These results suggested induction of type I IFN and p53-mediated DNA damage responses in the mutant cells, which was confirmed by gene set enrichment analysis (GSEA, Figure 3D). We also found upregulation of several ISGs in bone marrow and spleen of RH2^hKO^ animals by qRT-PCR (Figures S4A-B). In accordance with the ISG response on the transcriptome level, the type I IFN-induced surface protein Sca-1 was significantly upregulated on HSCs and PB lymphocytes from mutant compared to control mice as determined by flow cytometry (Figures 3E and S4C). Compatible with activation of a cGAS/STING response in RH2-deficient cells (Mackenzie et al., 2017), GSEA also revealed induction of NF-κB-dependent proinflammatory gene expression in the mutant cells.

**Figure 3.**
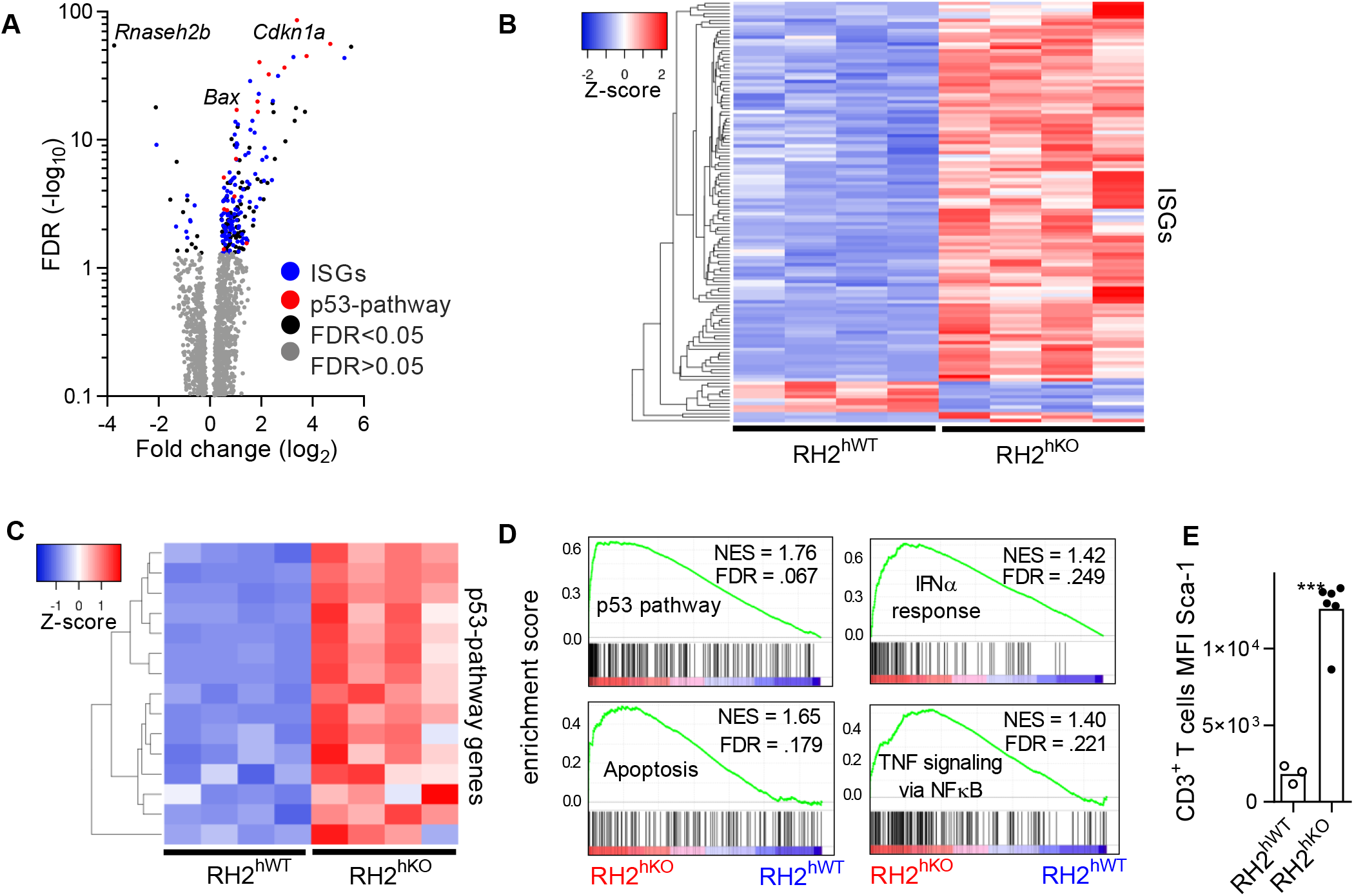
Genome damage in RH2^hKO^ mice activates p53 and type I IFN signaling. **A-D** Transcriptome analysis of BM granulocyte-macrophage progenitors (GMP, LE^-^K CD16/32^+^CD34^+^) isolated from RH2^hKO^ (n=4) and RH2^hWT^ control (n=4) mice revealed 218 differential expressed genes (false discovery rate (FDR) 5%). (A) Volcano plot (B) Differentially expressed interferon (IFN) stimulated genes (ISGs) and (C) p53 DNA repair pathway genes. (D) Gene set enrichment analysis showed upregulation of genes involved in p53, apoptosis, NFκB and IFNα signaling in RH2^hKO^ mice. **E** Sca-1 expression of PB T lymphocytes determined by FACS (n=3-6/genotype, individual mice and means are shown, unpaired Student’s t test).

### Attenuation of the p53 response rescues hematopoietic defects of RH2^hKO^ mice at the cost of accelerated leukemogenesis

We next investigated the impact of the activated p53 response on the phenotype of RH2^hKO^ mice and crossed the animals to *Trp53*^KO/KO^ mice (Donehower et al., 1992). Heterozygous loss of *Trp53* significantly ameliorated the hematopoietic deficits caused by absence of RH2 activity (Figure 4A). Erythrocyte counts were significantly increased in RH2^hKO^*Trp53*^KO/WT^ mice compared to the severe anemia of RH2^hKO^ controls. PB B cells, virtually absent in RH2^hKO^ animals, reached about 2/3 of control numbers, while PB T cells numbers were back to normal and the thrombocytosis was less pronounced in RH2^hKO^*Trp53*^KO/WT^ mice. Whereas BM was still hypocellular, HSCs and progenitor populations of RH2^hKO^*Trp53*^KO/WT^ animals were robustly increased in numbers compared to RH2^hKO^ BM, with MPPs and HPC-1s reaching control numbers (Figure 4B). Likewise, B and T cell development were partially rescued (Figures S5A-B). Competitive transplantation revealed that the additional loss of one *Trp53* allele partially restored the repopulation deficit of RH2-deficient HSCs upon primary (Figures 4 C-F) and secondary transplantation (Figures S5C-F). Heterozygous loss of p53 did not further increase the frequency of micronucleated erythrocytes in RH2-deficient hematopoiesis (Figure 4G). T lymphocytes (Figure 4H) and HSCs (Figure S5G) from RH2^hKO^*Trp53*^KO/WT^ mice still showed upregulation of the ISG Sca-1 compared to RH2^hWT^ controls.

**Figure 4.**
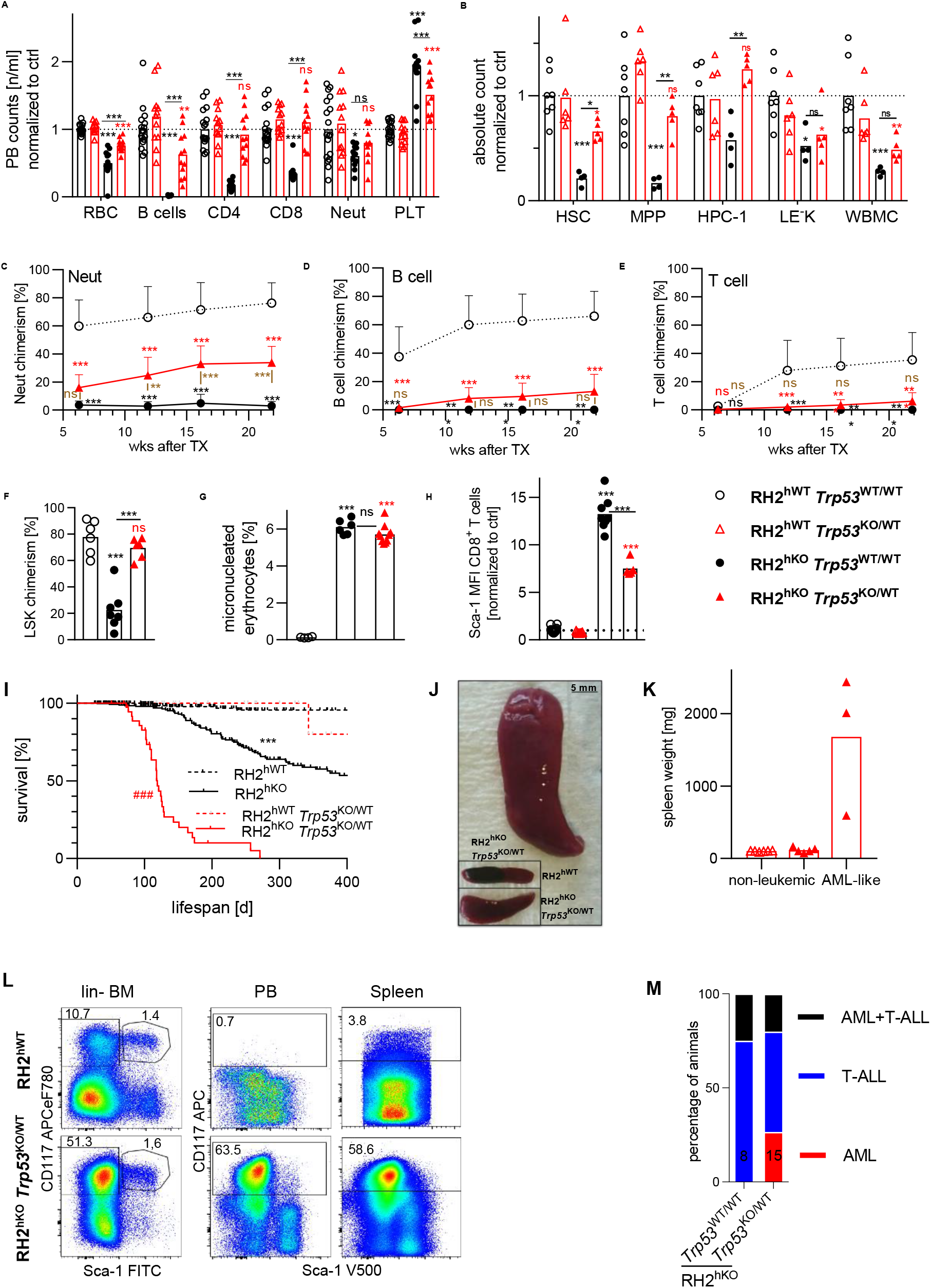
Attenuation of the p53 response rescues hematopoietic defects of RH2^hKO^ mice at the cost of accelerated leukemogenesis. **A-B** PB and BM of RH2^hKO^ mice with additional heterozygous loss of p53 (RH2^hKO^*Trp53*^KO/WT^, red closed triangles) were analyzed. Cell counts were normalized to RH2^hWT^*Trp53*^WT/WT^ controls (set to 1, open circles). Individual mice and means (bars) are shown, significance was calculated by 1way ANOVA with Holm-Sidak post test, RH2^hKO^*Trp53*^WT/WT^ (black asterisks) and RH2^hKO^*Trp53*^KO/WT^ (red asterisks) were compared to RH2^hWT^*Trp53*^WT/WT^ control mice, while significant differences between both RH2^hKO^ groups are underscored. PB cell counts (A) and absolute numbers of BM HSPC populations (B) isolated from 2 femora, 2 tibiae, and 2 pelvis are shown. **C-F** Competitive transplantation of HSCs isolated from either RH2^hKO^*Trp53*^KO/WT^, RH2^hKO^*Trp53*^WT/WT^ or RH2^hWT^*Trp53*^WT/WT^ mice. Each lethally-irradiated B6.CD45.1/CD45.2 recipient was transplanted with 200 donor HSCs (LEK CD48^-/lo^CD150^+^) previously mixed with 500,000 B6.CD45.1 BM competitor cells. PB neutrophil (Neut, C), B- (D) and T-cell (E) donor chimerism of primary recipients was determined (n=6-7 recipients/genotype, means and SD are shown, significance was calculated by repetitive measures 2way ANOVA with Holm-Sidak post test, comparison of RH2^hWT^ vs. RH2^hKO^*Trp53*^WT/WT^ black font, RH2^hWT^ vs. RH2^hKO^*Trp53*^KO/WT^ red font, RH2^hKO^*Trp53*^KO/WT^ vs. RH2^hKO^*Trp53*^WT/WT^ brown font). 22 weeks after transplantation, the donor chimerism of BM LSK cells was analyzed (F, individual mice and means (bars) are shown, statistics as in A, see Figures S5C-F for 2ndary transplantation). **G** The percentage of micronucleated erythrocytes was determined (display of data and statistics as in F). **H** Sca-1 median fluorescence intensity of CD8^+^ T cells from peripheral blood was analyzed (normalization, display and statistical analysis of data as in A-B). **I** Kaplan-Meier survival plot, significance was calculated by log-rank test (### = RH2^hKO^*Trp53*^WT/WT^ vs. RH2^hKO^*Trp53*^KO/WT^, ***= RH2^hKO^*Trp53*^WT/WT^ vs. RH2^hWT^*Trp53*^WT/WT^). **J** Splenomegaly in a RH2^hKO^*Trp53*^KO/WT^ mouse diagnosed with AML-like disease (bar = 5 mm, spleens from RH2^hWT^ and non-leukemic RH2^hKO^ *Trp53*^KO/WT^ animals are shown for comparison). **K** Spleen weights of control RH2^hWT^*Trp53*^KO/WT^ (n=6), non-leukemic RH2^hKO^*Trp53*^KO/WT^ (n=5) and AML-diseased RH2^hKO^*Trp53*^KO/WT^ animals (n=3, individual mice and means (bars) are shown). **L** Representative example of a leukemic RH2^hKO^*Trp53*^KO/WT^ mouse (lower row), which showed expansion of BM lin^-^CD117^hi^Sca-1^-^ cells (left), that disseminated to PB (middle) and spleen (right). Upper row depicts RH2^hWT^ control animal. **M** Distribution of disease entities in RH2^hKO^*Trp53*^WT/WT^ (n=8) and RH2^hKO^*Trp53*^KO/WT^ (n=13) diagnosed with leukemia.

While most parameters of RH2^hKO^ hematopoietic function were robustly improved by attenuated p53 expression, we observed a dramatically shortened median survival of RH2^hKO^*Trp53*^KO/WT^ animals in comparison to RH2^hKO^ animals (Figure 4I). In addition to development of T-ALL-like leukemia, some moribund RH2^hKO^*Trp53*^KO/WT^ animals featured grossly enlarged spleens with abnormal histology (Figures 4J-K, S6A). Flow cytometric analysis of BM, PB and spleen of these animals showed a massive expansion of aberrant lin^-^CD117^hi^Sca-1^-^CD201^-^CD34^-^CD150^-^CD48^-/lo^ myeloid progenitor cells (Figures 4L, S6B), thus a condition resembling acute myeloid leukemia (AML). Some mice developed both, T-ALL- and AML-like disease simultaneously (Figure 4M), which was revealed by gross enlargement of thymus and spleen by DP thymocytes (Figure S6C) and CD117^hi^Sca-1^-^ AML cells (Figure S6D), respectively, and both cell types coexisted in peripheral blood (Figure S6E).

We additionally generated few RH2^hKO^p53^KO/KO^ mice (n=4), which started to develop leukemic disease already by 8 weeks of age (Figure S6F). PB, BM and thymus phenotyping of these animals revealed a more pronounced rescue of hematopoietic defects than in RH2^hKO^*Trp53*^KO/WT^ mice (Figures S6G-J). Micronucleated erythrocytes were still elevated in RH2^hKO^*Trp53*^KO/KO^ animals, but slightly reduced in comparison to RH2^hKO^*Trp53*^WT/WT^ mice (Figure S6K), while the Sca-1 upregulation persisted (Figure S6L). In summary, attenuation of p53 signaling ameliorated the hematopoietic defects of RH2^hKO^ animals, but simultaneously fueled leukemia development.

### Signaling via the cGAS/STING axis has no impact on RER-deficient hematopoiesis

Loss of RER in hematopoietic cells led to DNA damage, micronucleus formation, and activation of innate immune pathways and, ultimately, malignant transformation. As genome damage and micronuclear DNA were shown to drive type I IFN via cGAS/STING in other models of RER deficiency (Mackenzie et al., 2016; Mackenzie et al., 2017), we crossed RH2^hKO^ mice to *Cgas*^KO/KO^, *Sting1*^GT/GT^ or *Ifnar1*^KO/KO^ animals. As expected, the enhanced expression of the type I IFN-induced surface protein Sca-1 in RH2^hKO^ mice was abrogated by each of these additional knock-outs (Figure 5A), demonstrating that the cGAS/STING axis relayed genome damage in RER-defective hematopoiesis into type I IFN signaling.

**Figure 5.**
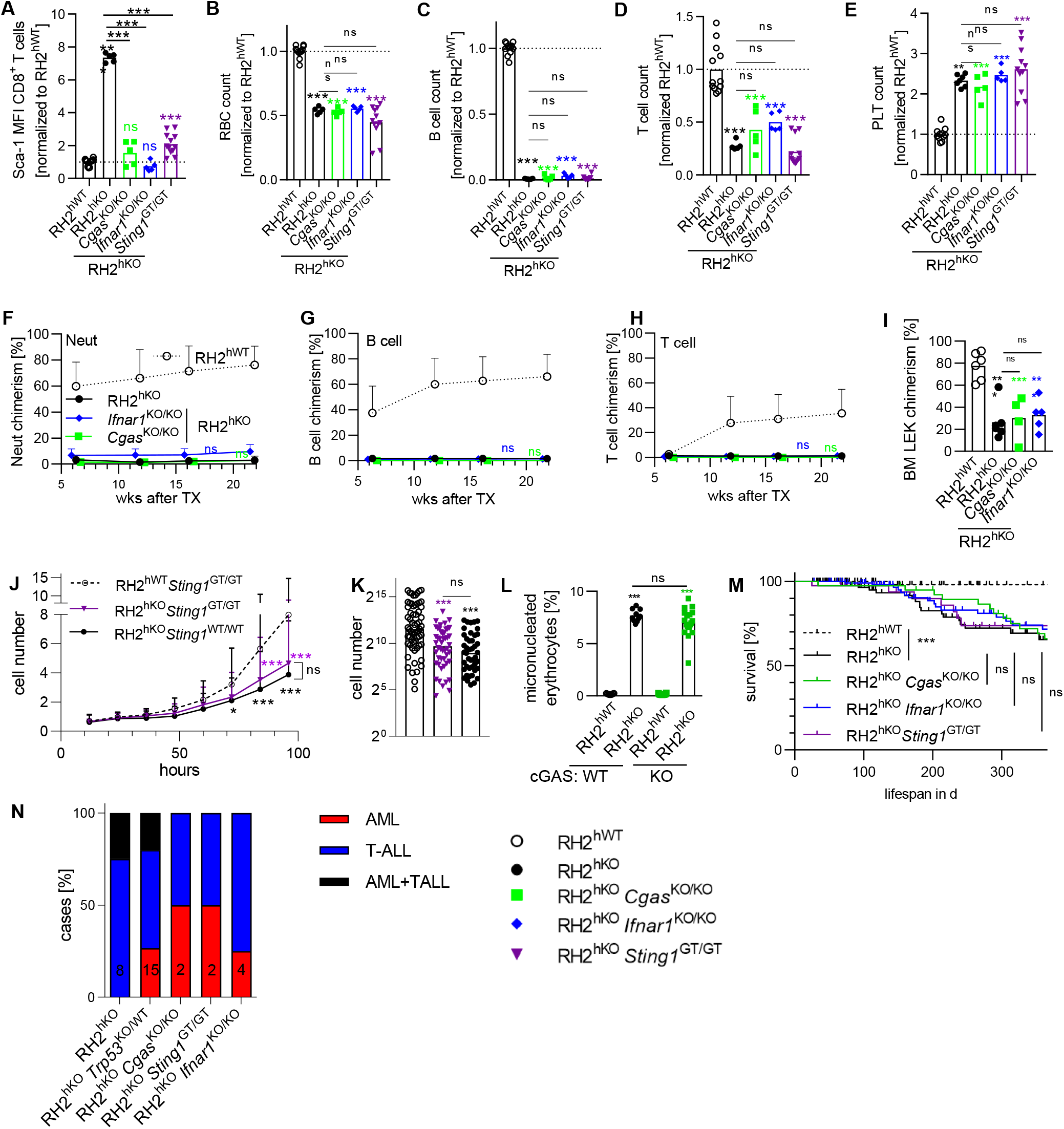
Signaling via the cGAS/STING axis has no impact on RER-deficient hematopoiesis. **A** Expression of the ISG Sca-1 was determined on PB CD8^+^ T cells isolated from RH2^hKO^ mice with an additional loss of either cGAS (*Cgas*^KO/KO^), IFNAR (*Ifnar1*^KO/KO^) or STING (*Sting1*^GT/GT^). RH2^hKO^ and RH2^hWT^ animals served as controls; data was normalized to the mean MFI of RH2^hWT^ mice, individual mice and means (bars) are shown, significance of either RH2^hWT^ (colored symbols) or RH2^hKO^ (underlined symbols) animals versus double KO mice was calculated by 1way ANOVA with Holm-Sidak post test. **B-E** Numbers of PB erythrocytes (B), B- (C), T-lymphocytes (D) and platelets (E). Display of data, normalization and statistics as in A. **F-I** Competitive transplantation of 200 purified HSCs (LEK CD48^-/lo^CD150^+^) isolated from RH2^hWT^, RH2^hKO^ or indicated double KO mice and injected together with 500,000 WT B6.CD45.1 BM competitor cells. The donor chimerism of PB neutrophils (Neut, F), B- (G) and T-cells (H) was determined (n=6-7 recipients/genotype, means and SD are shown, significance between RH2^hKO^ and double KO mice was calculated by RM 2way ANOVA with Holm-Sidak post test. (I) Chimerism of BM LEK cells 23 weeks after transplantation. Significance was calculated by 1way ANOVA with Holm-Sidak post test. **J-K** Single HSCs (LSK CD48^-/lo^CD150^+^CD34^-/lo^CD201^hi^, n=44-72/genotype & time point) were isolated from either RH2^hWT^*Sting1*^GT/GT^ (n=2), RH2^hKO^*Sting1*^WT/WT^ (n=3) or RH2^hKO^*Sting1*^GT/GT^ (n=3) mice and cultivated for 10 days. (J) Mean cell count and SD are shown, significance was calculated by repeated measures 2-way ANOVA with Holm-Sidak post test. (K) Colony size after 10 days of culture (median and individual colonies are shown, significance was calculated by 1way ANOVA with Holm-Sidak post test). **L** Percentage of micronucleated erythrocytes was determined (individual mice and means (bars) are shown, significance was calculated by 1way ANOVA with Holm-Sidak post test). **M** Kaplan-Meier survival curves of RH2^hKO^ mice with an additional loss of either cGAS, IFNAR or STING as well as RH2^hKO^ and RH2^hWT^ control animals (significance was calculated by log-rank test.) **N** Leukemia entities observed in RH2^hKO^ mouse strains with or without additional KO (number of diagnosed cases is given for each strain).

Activation of the cGAS/STING/IFN axis has antiproliferative effects and triggers cell death or senescence and potentially contributes to suppression of hematopoiesis, elimination of damaged cells and prevention of leukemia. Unexpectedly, however, we detected no effect of defective cGAS/STING or IFNAR signaling on blood cell production in RH2^hKO^ mice. All of the RH2^hKO^ double KO mouse strains developed macrocytic anemia, leukopenia and thrombocytosis at young age, indistinguishable from RH2 single KO mice (Figures 5B-E, S7A). Analysis of B lymphocyte and thymocyte development revealed persistent hematopoietic defects in RH2^hKO^*Sting1*^GT/GT^ mice (Figures S7B-C). The repopulation defect of transplanted RH2^hKO^ HSCs was not rescued by an additional loss of either cGAS or IFNAR (Figures 5F-I). Likewise, the proliferative potential of cultivated RH2^hKO^*Sting1*^GT/GT^ HSCs resembled that of RH2^hKO^ HSCs (Figures 5J-K). Abrogation of either cGAS, STING or IFNAR in RH2^hKO^ mice also had no effect on the frequency of micronucleated erythrocytes (Figures 5L, S7D-E). Collectively, in contrast to p53 signaling, the robust activation of cGAS/STING and type I IFN signaling in RH2^hKO^ mice is not a relevant factor impairing hematopoiesis.

In accordance with these findings, neither cGAS/STING nor type I IFN signaling contributed to prevention of malignant disease as RH2^hKO^ mice with an additional loss of either cGAS, STING or IFNAR exhibited a similar median survival as the RH2^hKO^ animals (Figure 5M). Analysis of few moribund RH2^hKO^ mice with additional loss of either cGAS, STING, or IFNAR revealed a similar pattern of leukemia (Figure 5N).

In summary, the cGAS/STING axis and type I IFN had no significant effect on the hematopoietic deficits or leukemogenesis of mice with severe chronic genome damage throughout the hematopoietic system.

### Loss of cGAS does not alter steady-state or stress hematopoiesis

Given that cGAS/STING signaling did not detectably impact on RER-deficient hematopoiesis, we next asked whether this pathway may be of relevance to control effects of spontaneous or acute DNA damage in mice with intact DNA repair. We first compared the size of hematopoietic cell populations between *Cgas*^KO/KO^ and WT control mice. However, we found no differences in hematopoietic stem and progenitor cell (HSPC) numbers or mature blood cell counts in control versus *Cgas*^KO/KO^ mice, neither young (Figure S8A-D) nor old (Figures S8E-F). In order to address whether cGAS deficiency affects the proliferative capacity of HSCs, we cultivated single HSCs purified from young and old *Cgas*^KO/KO^ or *Cgas*^WT/WT^ control mice and observed a similar growth pattern for both genotypes (Figures 6A-B, S8G-H). HSC functionality was also unchanged *in vivo* as demonstrated by repopulation experiments. Serial competitive transplantation of HSC purified from either *Cgas*^KO/KO^ or *Cgas*^WT/WT^ donor mice did not reveal any effect of cGAS deficiency on HSC repopulation capacity (Figures 6C-F, S8I-K).

**Figure 6.**
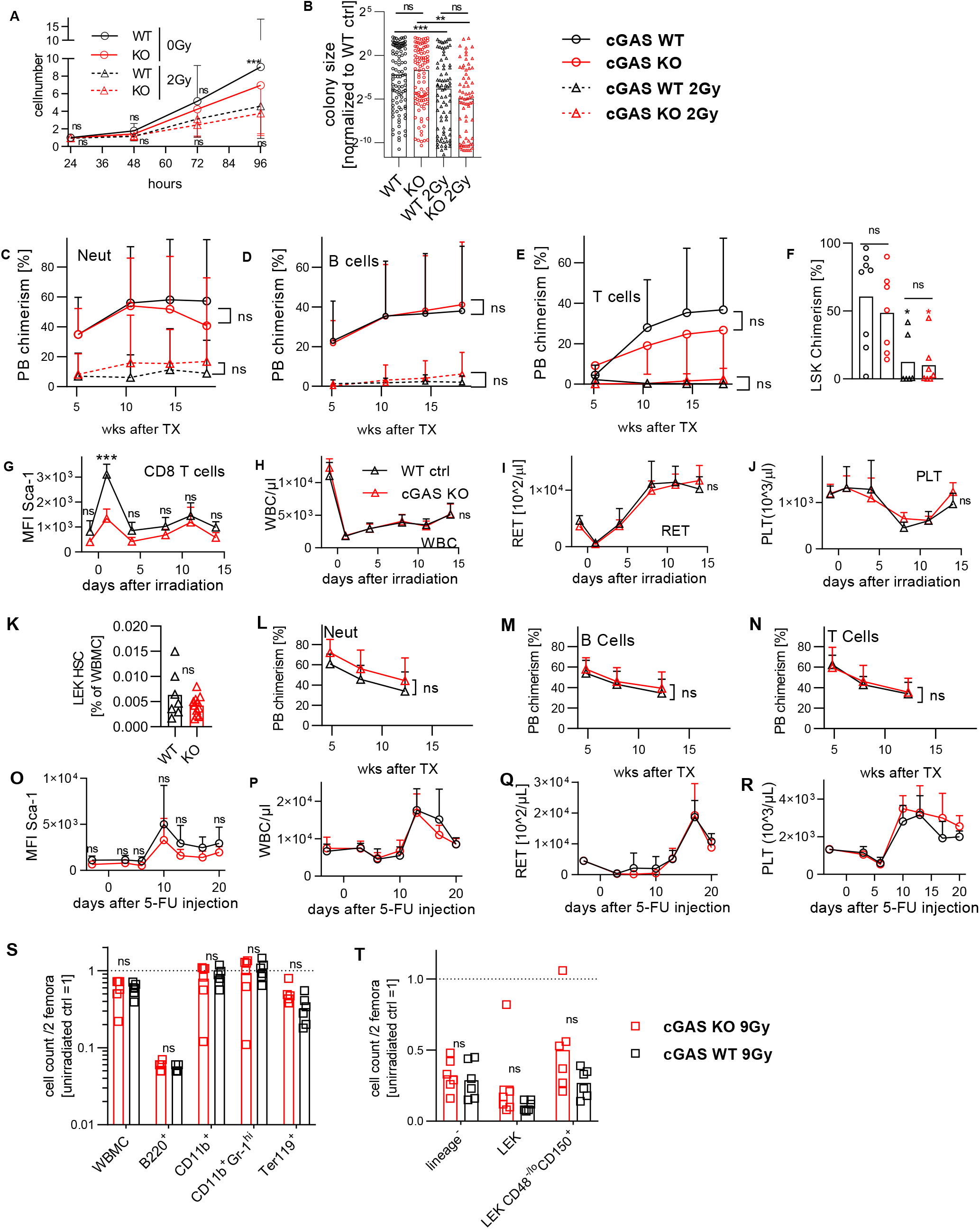
Loss of cGAS does not alter steady state or stress hematopoiesis. **A, B** Single HSCs (LSK CD48^-/lo^CD150^+^CD34^-/lo^CD201^hi^, n=70-103/condition) isolated from *Cgas*^KO/KO^ (red) or WT control (black) mice were cultivated for 10 days. A fraction of cells was exposed to 2 Gy γ-radiation (triangles) before culture. (A) Cells were counted for 96h every 24h (means and SD are shown, 2-way ANOVA with Sidak post test). (B) Colony size after 10 days of culture (medians and individual colonies are shown, significance was calculated by Mann-Whitney U test, pooled data from 2 individual experiments, mean colony size of cultivated HSCs from un-irradiated *Cgas*^WT/WT^ mice was set to 1). **C-F** 50 HSCs (LSK CD48^-/lo^CD150^+^ CD201^hi^) were isolated either from *Cgas*^KO/KO^ (red, n=5) or WT control (black, n=5) donor mice, mixed with 200,000 B6.CD45.1 WBMCs and competitively transplanted into lethally irradiated recipients (n=6-8/condition). Two additional recipient cohorts were transplanted with donor HSCs that received 2 Gy of γ-irradiation *ex vivo* (triangles). PB neutrophil (C), B- (D) and T- (E) lymphocyte chimerism of primary recipients was determined (means and SD are shown, significance was calculated by repeated measures 2way ANOVA). (F) Chimerism of BM LSK cells 18 weeks after transplantation (individual mice and means (bars) shown, significance was determined by 1way ANOVA with Sidak-Holm post test). **G-J** *Cgas*^KO/KO^ mice (red, n=6) and *Cgas*^WT/WT^ littermate controls (black, n=6) were exposed to 2 Gy γ-radiation. PB was repeatedly sampled, median Sca-1 fluorescence intensity of CD8^+^ T cells (G), leukocyte (H, WBC), reticulocyte (I, RET) and platelet (J, PLT) numbers were measured (means and SD shown, significance was calculated by repeated measures 2way ANOVA with Holm-Sidak post test). **K-N** *Cgas*^KO/KO^ mice (red, n=11) and *Cgas*^WT/WT^ littermate controls (black, n=7) were exposed to 2 Gy γ-radiation. (K) BM was isolated 30 days after irradiation and analyzed for HSC frequency (individual mice and means (bars) shown, significance was calculated by unpaired Student’s t test). To account for diminished HSC transplantation potential after irradiation, 10^7^ WBMCs from each mouse (either cGAS KO or WT) were mixed with 10^6^ B6.CD45.1 WBMCs and transplanted into a lethally irradiated B6.CD45.1/.2 recipient. PB chimerism of neutrophils (L), B- (M) and T (N) cells was analyzed (means and SD shown, significance was calculated by repeated measures 2way ANOVA with Holm-Sidak post test). **O-R** *Cgas*^KO/KO^ mice (red, n=6) and *Cgas*^WT/WT^ littermate controls (black, n=6) were injected with 5-FU. Display of data and statistics as in G-J. **S-T** *Cgas*^KO/KO^ mice (red, n=6) and *Cgas*^WT/WT^ littermate controls (black, n=6) were exposed to 9 Gy γ-radiation, sacrificed 10h later and BM was analyzed. Mature (S) and lineage-negative (T) BM populations were analyzed. Absolute cell counts from 2 femora were determined and normalized to un-irradiated control (set to 1, dotted line), individual mice and means (bars) are shown, significance was calculated by an unpaired Student’s t test.

To address the relevance of cGAS-mediated responses to acute DNA damage, we investigated the recovery of the hematopoietic system from a single exposure to 2 Gy whole body γ-radiation. This acute genotoxic stress indeed resulted in cGAS/STING activation, as reflected by a sharp increase in expression of the type I IFN-induced surface protein Sca-1 that was largely blunted in the *Cgas*^KO/KO^ animals (Figure 6G). As expected, irradiation caused a rapid loss of leukocytes, reticulocytes, and platelets (Figures 6H-J). After 5-10 days, white and red blood cell parameters started to recover, indicating compensatory mechanisms counteracting the radiation-induced blood cell loss. Neither the initial cell loss nor the following recovery phase was altered in *Cgas*^KO/KO^ animals compared to control mice. Likewise, the numbers of surviving HSPCs, B- and T-lymphocyte progenitors (Figures S8L-O) after irradiation were not changed by lack of cGAS. Competitive transplantation of WBMCs isolated from *Cgas*^KO/KO^ and WT control animals 30 days after 2 Gy whole body γ-irradiation confirmed that survival and regeneration of HSCs was not influenced by cGAS deficiency (Figure 6K-N). Moreover, *in vitro*-proliferation of sorted HSCs irradiated with 2 Gy prior to cultivation was not altered by the absence of cGAS (Figures 6A-B, S8G-H), and also repopulation potential of γ-irradiated HSCs, reduced compared to unirradiated HSCs as expected, was not affected by cGAS deficiency (Figures 6C-F). In addition to γ-irradiation, we also induced acute replication stress in cGAS-deficient and control mice by exposure to the myeloablative drug 5-fluorouracil (5-FU). This stressor causes rapid loss of cycling stem and progenitor cells and surviving cells subsequently initiate hematopoietic regeneration. However, loss of cGAS had no impact on the hematopoietic recovery after 5-FU-induced myeloablation (Figures 6O-R, S8P-S).

A previous study (Jiang et al., 2019) reported that cGAS suppresses homologous recombination in a STING-independent fashion, and that loss of cGAS accordingly enhanced the resistance of hematopoietic cells to ionizing radiation. As we had not observed cGAS-mediated effects on the loss of hematopoietic cells upon whole body irradiation with a dose of 2 Gy, we repeated this experiment with the same dose and observation interval as in the study of Jiang et al. However, cGAS deficiency did not protect BM cells from radiation-induced cell death evidenced by massive cell loss and similar numbers of HSPCs as well as mature blood cells in both genotypes 10 hours after 9 Gy whole body γ-irradiation (Figures 6S-T, S8T).

In summary, loss of cGAS had no effect on steady-state hematopoiesis or stem cell function in young or old mice. *In vivo*, cGAS appeared irrelevant for the survival, function and recovery of hematopoietic stem and progenitor cells after acute genotoxic insults.

## Discussion

We report a genetic model of chronic genome damage due to defective DNA repair, causing massive cytopenia and leukemia, and in which both, p53 and cGAS/STING pathways are robustly activated. While p53 signaling eliminates large numbers of damaged cells and efficiently delays onset of leukemia, cGAS/STING signaling appears irrelevant for hematopoietic function and prevention of malignant transformation. In mice with intact DNA repair, we found that cGAS deficiency does not alter the response of the hematopoietic system to spontaneous or acute DNA damage.

The hematopoietic system is the most regenerative mammalian tissue and accounts for ∼85% of the daily cellular turnover (Cosgrove et al., 2021; Sender and Milo, 2021). Genome damage resulting from loss of RER and subsequent DSB formation primarily affects replicating cells (Cerritelli and Crouch, 2016; Kellner and Luke, 2020; Williams et al., 2016). Accordingly, erythropoiesis, which accounts for the main share of blood cell production, was strongly affected. However, lymphopoiesis was even more severely suppressed, suggesting that DNA damage related to loss of RER is either more strictly controlled in lymphocytes, e.g. by induction of cell death, or that lymphocytes favor error-prone repair by NHEJ (Libri et al., 2022) and thereby massively accumulate fatal genome damage. Interestingly, RH2^hKO^ mice developed thrombocythemia, while production of all other blood cell lineages was impaired. This seemingly contradictory finding is likely explained by the observation that large numbers of thrombocytes can be generated by direct differentiation of HSCs into megakaryocytes, a process that requires only minimal cell division (Morcos et al., 2022), while supply of all other mature blood cells heavily relies on cell division of stem and progenitors cells.

Additional loss of p53 significantly rescued HSC function and blood cell generation in RH2^hKO^ mice, but also accelerated leukemogenesis, demonstrating that p53 is crucial for orchestrating the response to genome damage ensuing from RER-deficiency. Similar observations were made in neurons (Aditi et al., 2021), skin (Hiller et al., 2018) and gut (Aden et al., 2019) epithelium. Interestingly, the thrombocythemia of RH2^hKO^ mice was reversed by loss of p53, in line with reported roles for p53 signaling in regulating megakaryocyte endoreplication and platelet numbers (Apostolidis et al., 2012; Yang et al., 2020).

In addition to the p53-mediated DNA damage response, cGAS/STING signaling was robustly activated by RER deficiency and resulted in massive production of type I IFN and TNF. However, inactivation of either cGAS, STING or IFNAR signaling abrogated the cytokine responses but did not rescue anemia or lymphopenia of RER-deficient mice. This was unexpected as STING activation was shown to induce cell death (Gaidt et al., 2017; Gulen et al., 2017; Hayman et al., 2021; Paludan et al., 2019) and to trigger senescence (Glück et al., 2017; Yang et al., 2017) and is regarded as an anti-oncogenic principle counteracting malignant transformation by removing or inactivating excessively genome damage cells (Li and Chen, 2018). However, we found that neither survival nor incidence of leukemia of RH2^hKO^ mice was affected by additional inactivation of cGAS, STING or IFNAR. Similar observations were made in a mouse model of neuronal RH2 deficiency, in which loss of cGAS dampened inflammatory signaling, but did not rescue neuropathology (Aditi et al., 2021). Moreover, analysis of RH2-deficient astrocytes derived from Aicardi-Goutières syndrome patients suggested that neurotoxicity developed independently of type I IFN signaling (Giordano et al., 2022). cGAS was recently identified as a pre-dominantly nuclear protein, that binds to chromatin but its catalytic activity is inhibited by nucleosomes (Volkman et al., 2019; Zierhut et al., 2019). Nuclear cGAS was reported to maintain genome integrity by stabilizing replication forks (Chen et al., 2020) and may explain that presence of cGAS protected from colitis-associated colon cancer through a STING-independent mechanism (Hu et al., 2021). In contrast, two studies claimed that nuclear cGAS suppresses DNA DSB repair by homologous recombination in a STING-independent manner (Jiang et al., 2019; Liu et al., 2018), thereby sensitizing cells to programmed cells death after acute genotoxic stress as well as promoting genome instability and tumorigenesis. Accordingly, Jiang et al. reported markedly improved survival of cGAS-deficient BM cells upon ionizing radiation. However, in our models of either chronic or acute DNA damage, cGAS-deficiency did not improve survival or regeneration of hematopoietic cells. We conclude that, in a RER-deficient hematopoietic system, cGAS does neither prevent nor promote DNA damage accumulation and leukemia development. This is in line with mechanisms that selectively silence cGAS in HSCs (Xia et al., 2018) and with reports of HSCs being refractory to massive type I IFN stimulation and thereby protected from over-activation and exhaustion (Pietras et al., 2014; Wu et al., 2018).

Collectively, we show that loss of RER in the hematopoietic system caused profound defects in blood cell development and predisposed to leukemia. p53 activation in RER-deficient hematopoiesis resulted in impaired blood formation but also efficiently delayed leukemia development. In contrast, cGAS/STING signaling did not alter the capacity of the hematopoietic system to cope with acute or chronic DNA damage. Future work will clarify whether cGAS/STING responses may represent a back-up surveillance pathway that becomes relevant upon loss of p53 function, as occurring frequently in malignant cells.

## Supporting information

Tables S1-3 GMP transcriptome analysis

Table S4 Antibodies

Table S5 Primers for qRT-PCR

## Acknowledgements

The authors thank Christa Haase, Livia Schulze and Madeleine Rickauer for expert technical assistance. This study was funded by the Fritz-Thyssen-Stiftung (Az. 10.19.1.013MN to A.G.), Deutsche Forschungsgemeinschaft (DFG, German Research Foundation) – Project-ID 369799452 – TRR237 Nucleic Acid Immunity, project B17 to A.R., project B19 to R.B. and project B18 to A.R.-W. L.N. was supported by the Elke-Kröner-Promotionskolleg.

## Author Contributions

A. G., R.B. and A.R. conceptualized the study. Experimental work and data analysis was carried out by N.D. and L.N. with assistance from C.M.M., S.C.R., M.N.F.M, A.L. and B.H. H.L. and A.R.-W. discussed data. RNA-sequencing and transcriptome analysis was performed by A.D., M.L. and R.B. A.G., A.R. and R.B. wrote the manuscript. All authors read, commented, and agreed on the final version of the manuscript.

## Conflict of Interest

The authors declare no conflict of interests

## Figure legends

**Figure S1.**
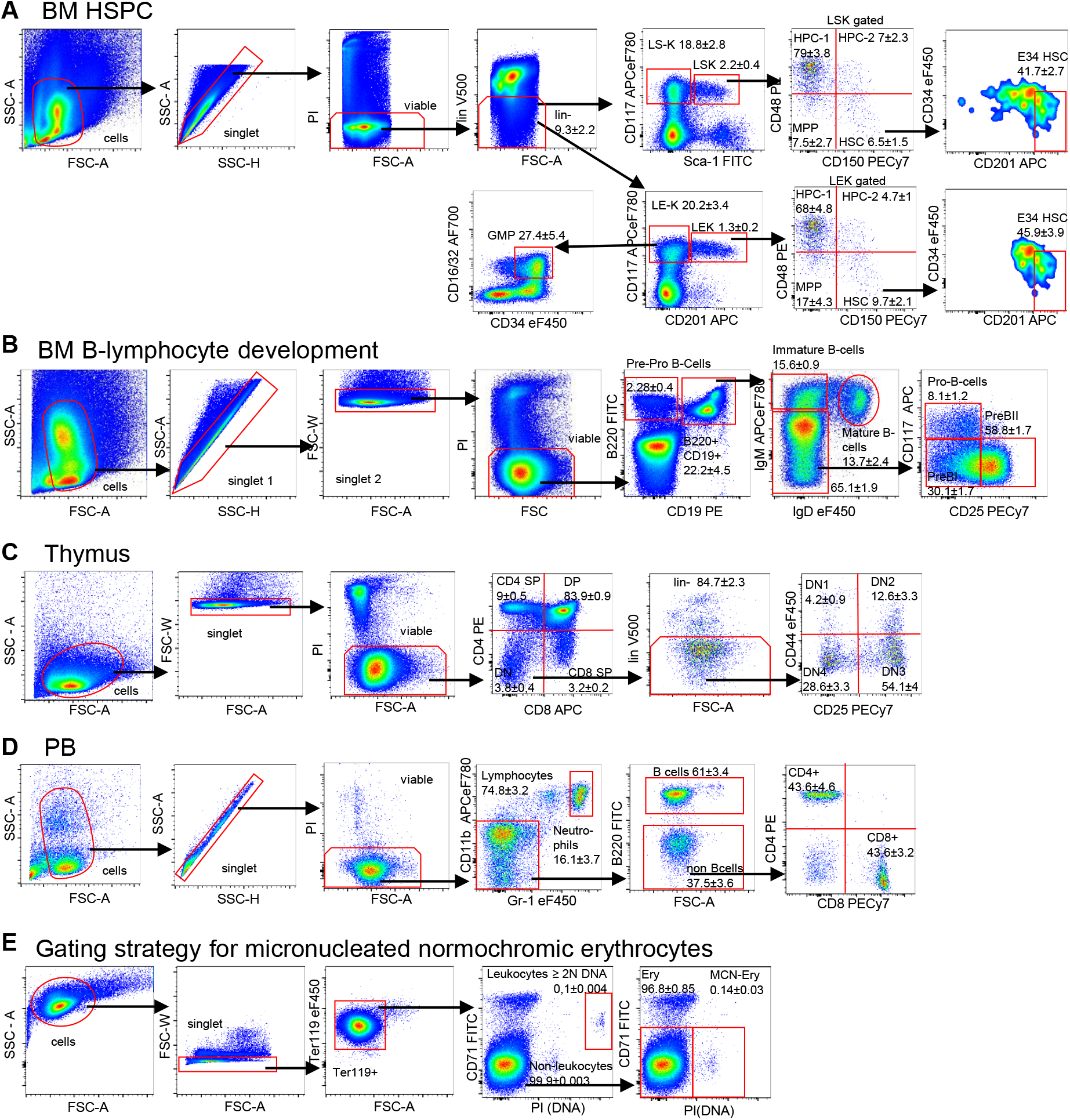
Gating strategies for flow cytometry. **A** Representative gating strategy for identification of bone marrow hematopoietic stem and progenitor populations. RNase H2-deficient mouse strain spontaneously induce a type I interferone signature and since Sca-1 represents a type I interferone-stimulated gene (ISG), we used an alternative gating strategy employing CD201 (EPCR) for separation of early hematopoietic stem and progenitor cells (Sca-1^+^/CD201^+^) and committed hematopoietic progenitors (Sca-1^-^/CD201^-^). (*LS*^*-*^*K*, lin^-^ Sca-1^-^ CD117^+^ cells; *LSK*, lin^-^ Sca-1^+^ CD117^+^ cells; *restricted hematopoietic progenitor 1 (HPC-1)*, LSK CD48^hi^ CD150^-^; *restricted hematopoietic progenitor 2 (HPC-2)*, LSK CD48^hi^ CD150^+^; *multipotent progenitor (MPP)*, LSK CD48^-/lo^ CD150^-^; *hematopoietic stem cell (HSC)*, LSK CD48^-/lo^ CD150^+^; *E34 HSC*, LSK CD48^-/lo^ CD150^+^ CD34^-/lo^ CD201^hi^; *LE*^*-*^*K*, lin^-^ CD201^-^ CD117^+^ cells; *LEK*, lin^-^ CD201^+^ CD117^+^ cells; *restricted hematopoietic progenitor 1 (HPC-1)*, LEK CD48^hi^ CD150^-^; *restricted hematopoietic progenitor 2 (HPC-2)*, LEK CD48^hi^ CD150^+^; *multipotent progenitor (MPP)*, LEK CD48^-/lo^ CD150^-^; *hematopoietic stem cell (HSC)*, LEK CD48^-/lo^ CD150^+^; *E34 HSC*, LEK CD48^-/lo^ CD150^+^ CD34^-/lo^ CD201^hi^; *granulocyte-macrophage progenitor (GMP)*, LE^-^K CD16/32^+^ CD34^+^; percentage among parent population ± SD is given and was calculated from RH2^hWT^ mice (n=7)). For lineage staining biotin conjugated primary antibodies against murine CD11b, CD8, CD19, NK1.1, Gr-1, CD4, CD3, Ter119 and B220 were used and followed by staining with Streptavidin conjugated to BD Horizon V500. **B** Representative gating strategy for identification of bone marrow B-lymphocyte populations (*Pre-Pro B-Cells*, CD19^-^ B220^+^; *Immature B-Cells*, CD19^+^ B220^+^ IgM^+^ IgD^-^; *Mature B-Cells*, CD19^+^ B220^+^ IgM^+^ IgD^+^; *Pro-B-Cells*, CD19^+^ B220^+^ IgM^-^ IgD^-^ CD117^+^ CD25^-^; *PreBI*, CD19^+^ B220^+^ IgM^-^ IgD^-^ CD117^-^ CD25^-^; *PreBII*, CD19^+^ B220^+^ IgM^-^ IgD^-^ CD117^-^ CD25^+^; percentage of parent population ± SD is given and was calculated from RH2^hWT^ mice (=7)). **C** Representative gating strategy for identification of thymocyte populations (*CD4 single-positive (SP) thymocytes*, CD4^+^ CD8^-^; *CD8 SP thymocytes*, CD4^-^ CD8^+^; *CD4/8*-*double positive (DP) thymocytes*, CD4^+^ CD8^*+*^; *CD4/8*-*double negative (DN) thymocytes*, CD4^-^ CD8^-^ ; *DN1*, CD4^-^ CD8^-^ lin^-^ CD25^-^ CD44^+^; *DN2*, CD4^-^ CD8^-^ lin^-^ CD25^+^ CD44^+^; *DN3*, CD4^-^ CD8^-^ lin^-^ CD25^+^ CD44^-^; *DN4*, CD4^-^ CD8^-^ lin^-^ CD25^-^ CD44^-^); percentage of parent populations ± SD was calculated from RH2^hWT^ mice (n=7)). Lineage staining against B220, NK1.1, CD11b, Gr-1, Ter119, CD19, CD49b surface markers was performed using biotin conjugated antibodies and Streptavidin BD Horizon V500 as secondary reagent. **D** Representative gating strategy for identification of peripheral blood (PB) leukocytes (*neutrophils (Neut)*, CD11b^+^ Gr1^hi^; *B-cells*, CD11b^-^ GR1^-^ B220^+^; *CD4*^*+*^ *T-cells*, CD11b^-^ GR1^-^ B220^-^ CD4^+^ CD8^-^ and *CD8*^*+*^ *T-cells*, CD11b^-^ GR1^-^ B220^-^ CD4^-^ CD8^+^; percentage of parent populations ± SD is given and was calculated from RH2^hWT^ mice (n=7)). **E** Representative gating strategy for identification of micronucleated (MCN) erythrocytes (Ery). PB was fixed using -80°C methanol for at least 24h prior to staining. Fixed erythrocytes were digested using RNase A and stained with Ter119, CD71 and propidium iodide (PI). Nucleated leukocytes (≥2N DNA content) and CD71^+^ reticulocytes were discriminated and normochromic (CD71^-^) erythrocytes were analysed for micronuclei (PI^+^, percentage of parent populations ± SD is given and was calculated from RH2^hWT^ mice (n=4)).

**Figure S2.**
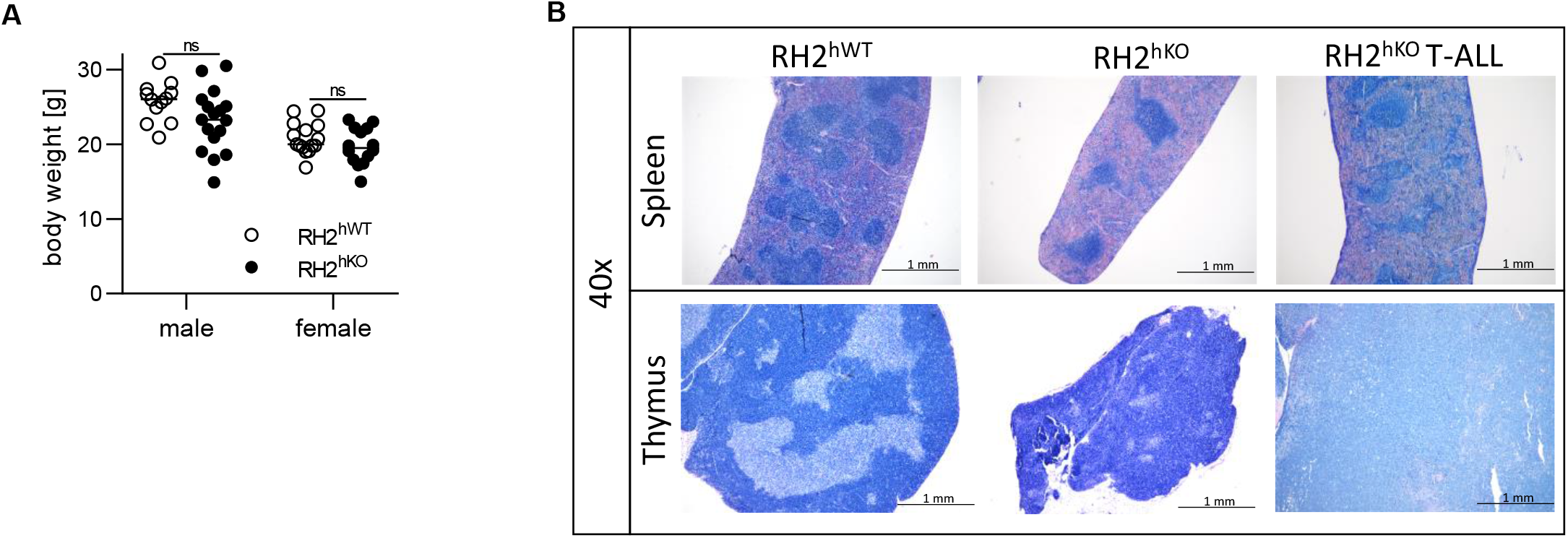
Hematopoietic loss of RER results in genome instability and predisposes to leukemia. **A** Body weight of age- and sex-matched, non-leukemic RH2^hKO^ (closed circles) and RH2^hWT^ (open circles) mice (aged 6-16 wks, n=12-17/group). Significance was calculated by an unpaired student’s t-test. **B** Representative micrographs of spleen (upper row) and thymus (lower row) tissue isolated from RH2^hWT^ (left), non-leukemic RH2^hKO^ (middle) and leukemic RH2^hKO^ (right) animals. Sections were stained by May-Grünwald-Giemsa.

**Figure S3.**
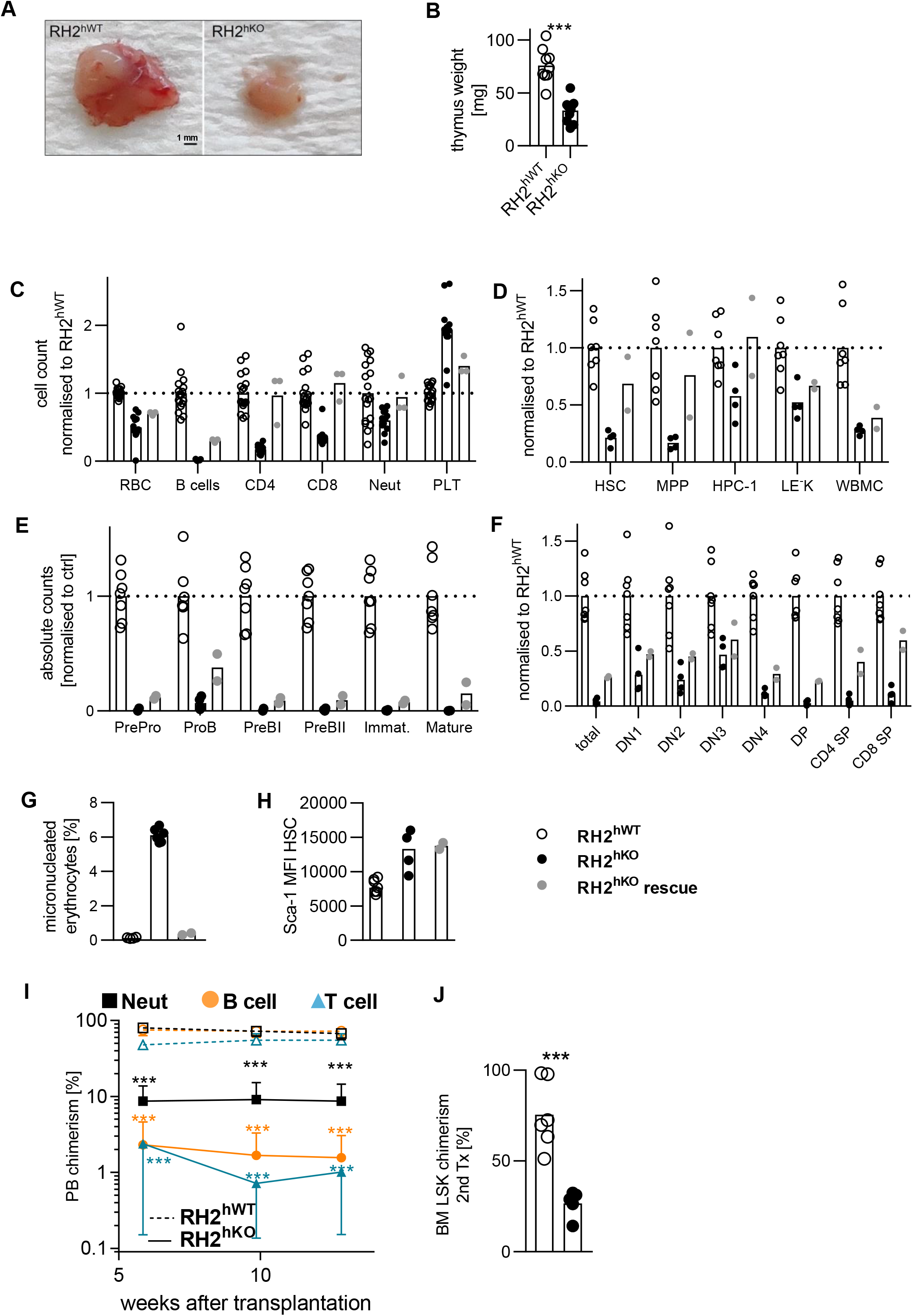
Chronic genome damage impairs blood cell production and HSC fitness. **A** Representative example of thymi isolated from a non-leukemic RH2^hKO^ (right) or RH2^hWT^ (left) mouse (scale bar 1 mm). **B** Weight of thymi isolated from non-leukemic RH2^hKO^ mice and RH2^hWT^ littermates (n=9-10/genotype, age 8-16 wks, individuals and means (bars) are shown, significance was calculated by an un-paired Student’s t test). **C-H** Few RH2^hKO^ mice exhibited ameliorated cytopenia and were designated as “RH2^hKO^ rescue” (grey circles). PB (C), HSPC (D), B lymphocyte (E) and thymocyte (F) counts; same RH2^hKO^ and RH2^hWT^ animals, display and normalization of data as in Figures 2A-D. (G) Percentages of micronucleated erythrocytes (H) Median fluorescence intensity (MFI) of Sca-1 surface expression among HSCs (LEK CD48^-/lo^ CD150^+^). **I-J** 5×10^6^ WBMCs were isolated from primary recipients of either RH2^hKO^ or RH2^hWT^ HSCs (experiment shown in Figures 2H-I) and serially transplanted into lethally irradiated secondary recipients (n=6/genotype). PB (I) and BM LSK (J) chimerism were analyzed (display of data and statistics as in Figures 2H-I).

**Figure S4.**
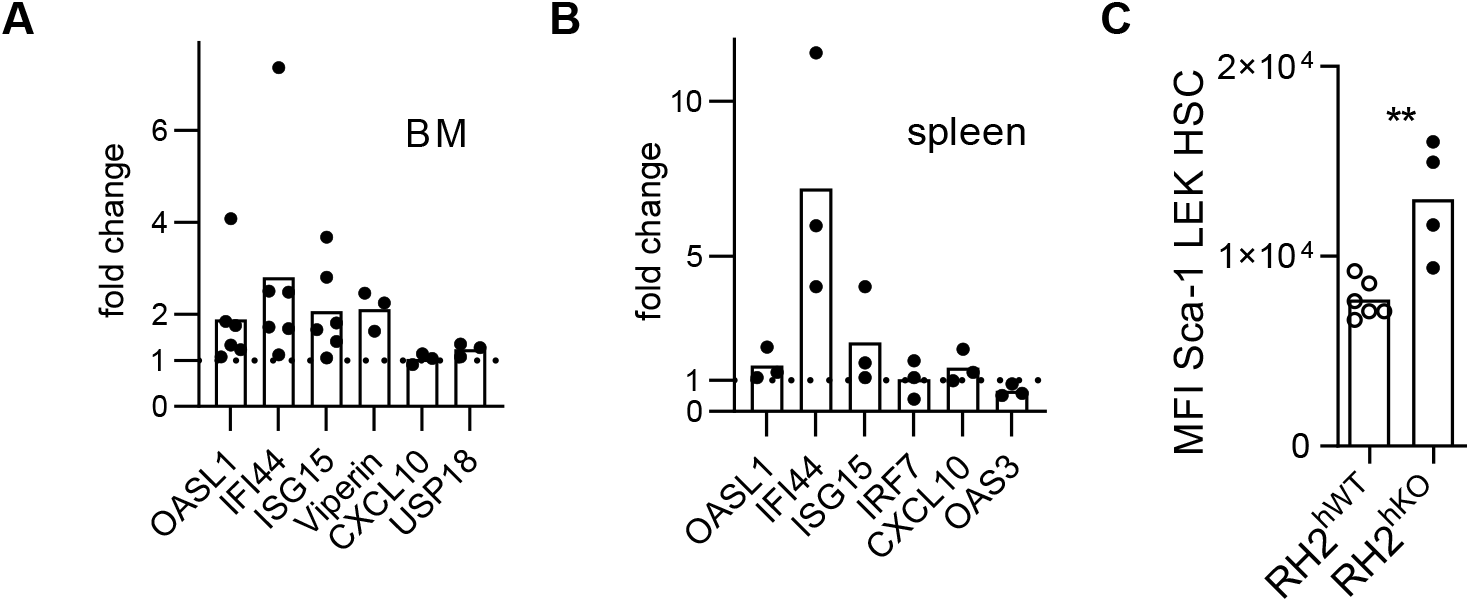
Genome damage in RH2^hKO^ mice activates p53 and type I IFN signaling. **A-B** Expression of ISGs was determined by quantitative reverse transcription (qRT)-PCR in BM cells (A) and splenocytes (B) isolated from RH2^hKO^ (individual mice and means (bars) are shown) and RH2^hWT^ control mice (dotted line). **C** Surface Sca-1 expression was determined on HSCs (LEK CD48^-/lo^ CD150^+^, significance was calculated by unpaired Student’s t test).

**Figure S5.**
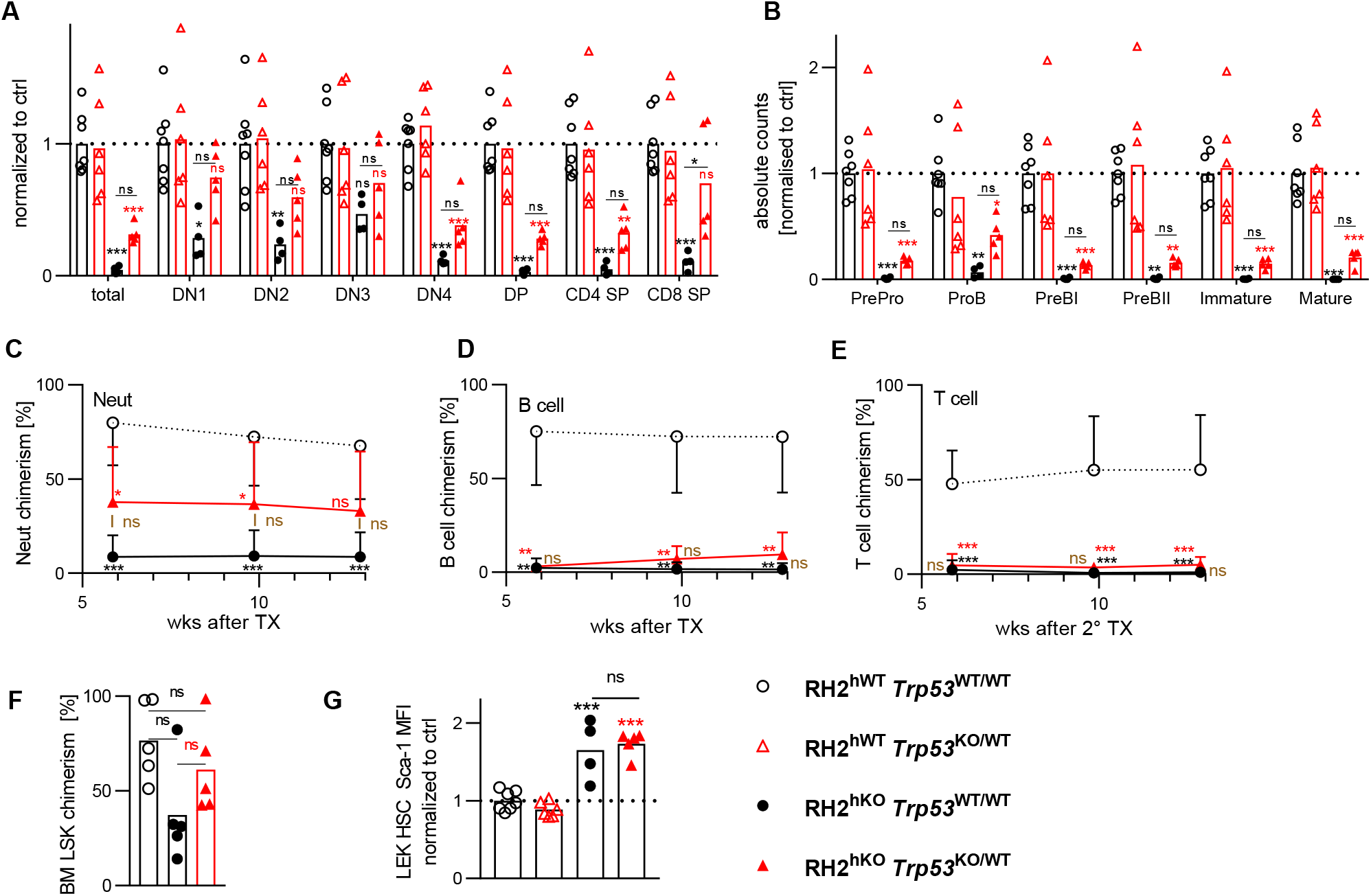
Attenuation of the p53 response rescues the hematopoietic defects of RH2^hKO^ mice. **A-B** Thymocyte (A) and BM B cell (B) development (display and normalization of data and statistics as in Figures 4A-B). **C-F** 5 × 10^6^ whole bone marrow cells isolated from each primary recipients (Figures 4C-F) were transplanted into a sex-matched secondary recipients. Chimerism of PB neutrophils (C), B- (D) and T- (E) cells (display of data and statistics as in Figures 4C-E) and BM LSK cells (F, 14 weeks after secondary transplantation, individual mice and means (bars) shown, 1way ANOVA with Holm-Sidak post test) was analyzed. **G** Sca-1 MFI of HSCs (LEK CD48^-/lo^CD150^+^) was calculated (individuals and means (bars) shown, significance was calculated by 1way ANOVA with Holm-Sidak post test

**Figure S6.**
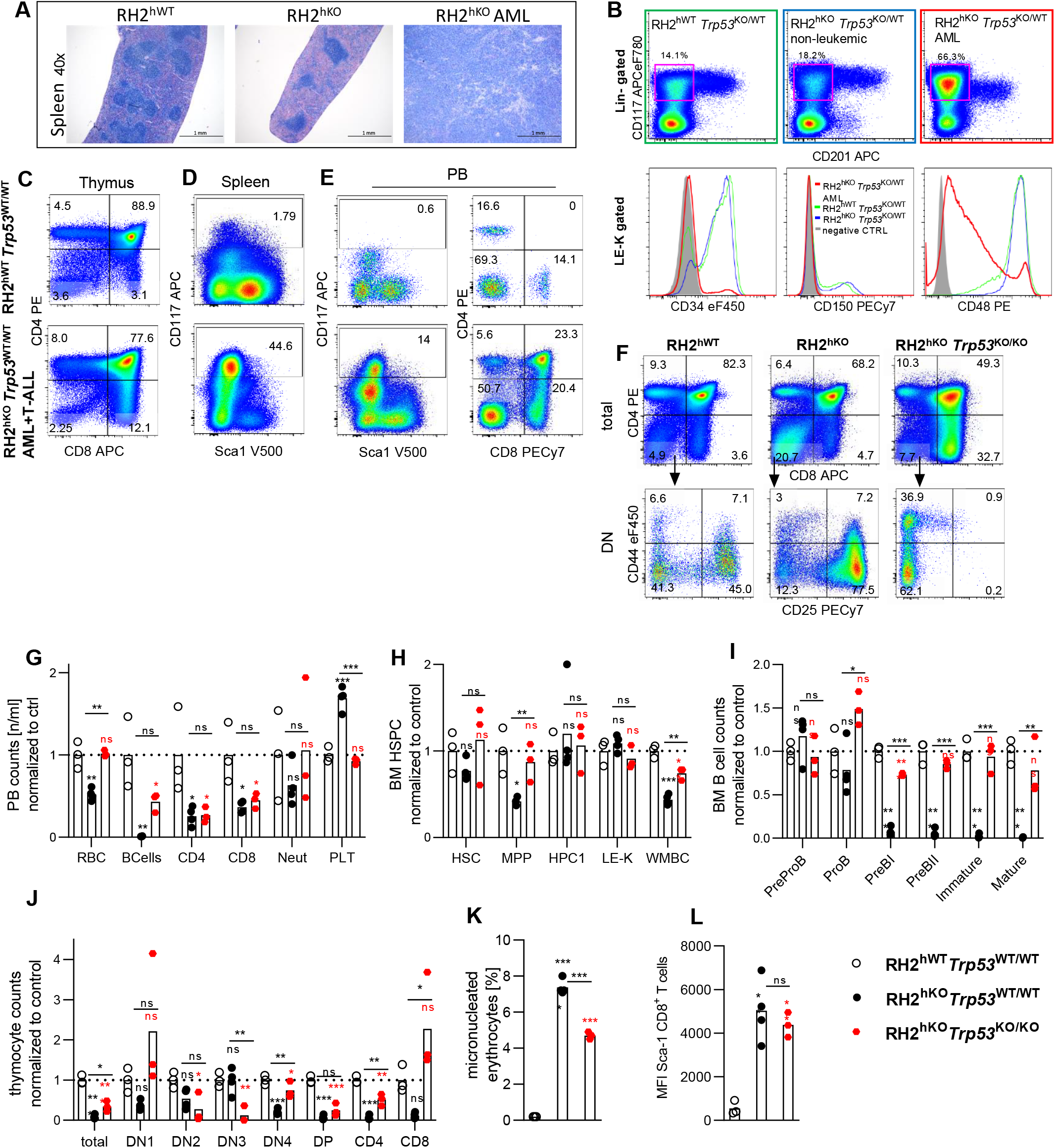
Loss of *Trp53* accelerates leukemogenesis of RH2^hKO^ mice. **A** Spleen histology (May-Grünwald Giemsa staining) from a representative RH2^hKO^*Trp53*^KO/WT^ mouse diagnosed with AML-like disease and control animals. **B** Immuno-phenotypical characterization of BM cells isolated from a representative RH2^hKO^*Trp53*^KO/WT^ mouse diagnosed with AML and non-leukemic controls (upper row shows lineage-negative BM cells, lower row shows histograms of lineage^-^CD201^-^Kit^hi^ (LE^-^K) cells. Fluorescence minus-one controls served as negative controls). **C-E** Representative example of an RH2^hKO^*Trp53*^WT/WT^ mouse (lower row) which developed AML- and T-ALL-like disease simultaneously. Shown are thymus (C), Spleen (D) and peripheral blood (E). Upper row depicts RH2^hWT^ control animal. **F-L** Phenotypical characterization of RH2^hKO^ mice with homozygous p53 deficiency. All RH2^hKO^*Trp53*^KO/KO^ animals (n=4) developed leukemia by 8 weeks of age. (F) Representative dot plots of total (upper row) and DN (lower row) thymocyte suspensions isolated from RH2^hWT^, RH2^hKO^, and RH2^hKO^*Trp53*^KO/KO^ animals. (G-L) RH2^hWT^*Trp53*^WT/WT^ (n=3), RH2^hKO^*Trp53*^WT/WT^ (n=4) and RH2^hKO^*Trp53*^KO/KO^ (n=3) mice (8 wks of age) were analyzed by flow cytometry. PB (G), BM HSPC (H), BM B lymphocyte (I) and thymocyte (J) development were analyzed. Individual mice and means (bars) are shown, data was normalized to RH2^hWT^ control (set to 1), significance was calculated using 1way Anova with Holm-Sidak post test. (K) Micronucleated erythrocytes and (L) Sca-1 expression of CD8+ T-cells (significance was calculated by 1way ANOVA with Holm-Sidak post test).

**Figure S7.**
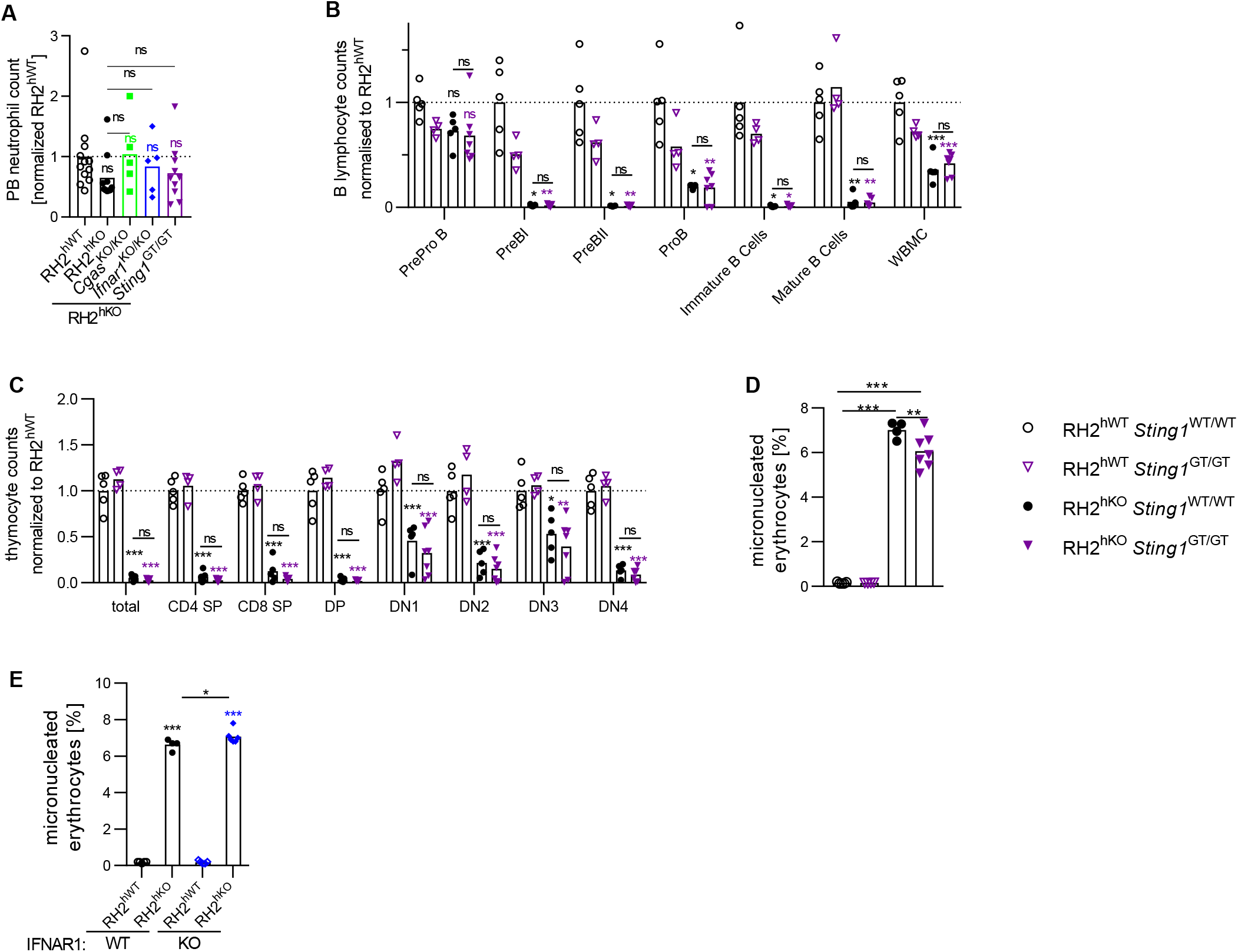
Signaling via the cGAS/STING axis has no impact on RER-deficient hematopoiesis. **A** Numbers of PB neutrophils. Normalization, display of data and statistics as in Figure 6A. **B-C** BM B lymphocyte (B) and thymocyte (C) development of RH2^hKO^ mice with additional loss of STING (RH2^hKO^*Sting1*^GT/GT^) were analyzed. Cell counts were normalized to RH2^hWT^*Sting1*^WT/WT^ controls (set to 1, dotted line). Individual mice and means (bars) are shown, significance was calculated by 1way ANOVA with Holm-Sidak post test. **D** The frequencies of micronucleated erythrocytes from PB of RH2^hWT^, RH2^hWT^*Sting1*^GT/GT^, RH2^hKO^ and RH2^hKO^*Sting1*^GT/GT^ mice were analyzed (individuals and means (bars) are shown, significance was calculated by 1way ANOVA with Holm-Sidak post test). **E** The frequencies of micronucleated erythrocytes from PB of RH2^hWT^, RH2^hWT^*Ifnar1*^KO/KO^, RH2^hKO^ and RH2^hKO^*Ifnar1*^KO/KO^ were determined (individuals and means (bars) are shown, significance was calculated by 1way ANOVA with Holm-Sidak post test).

**Figure S8.**
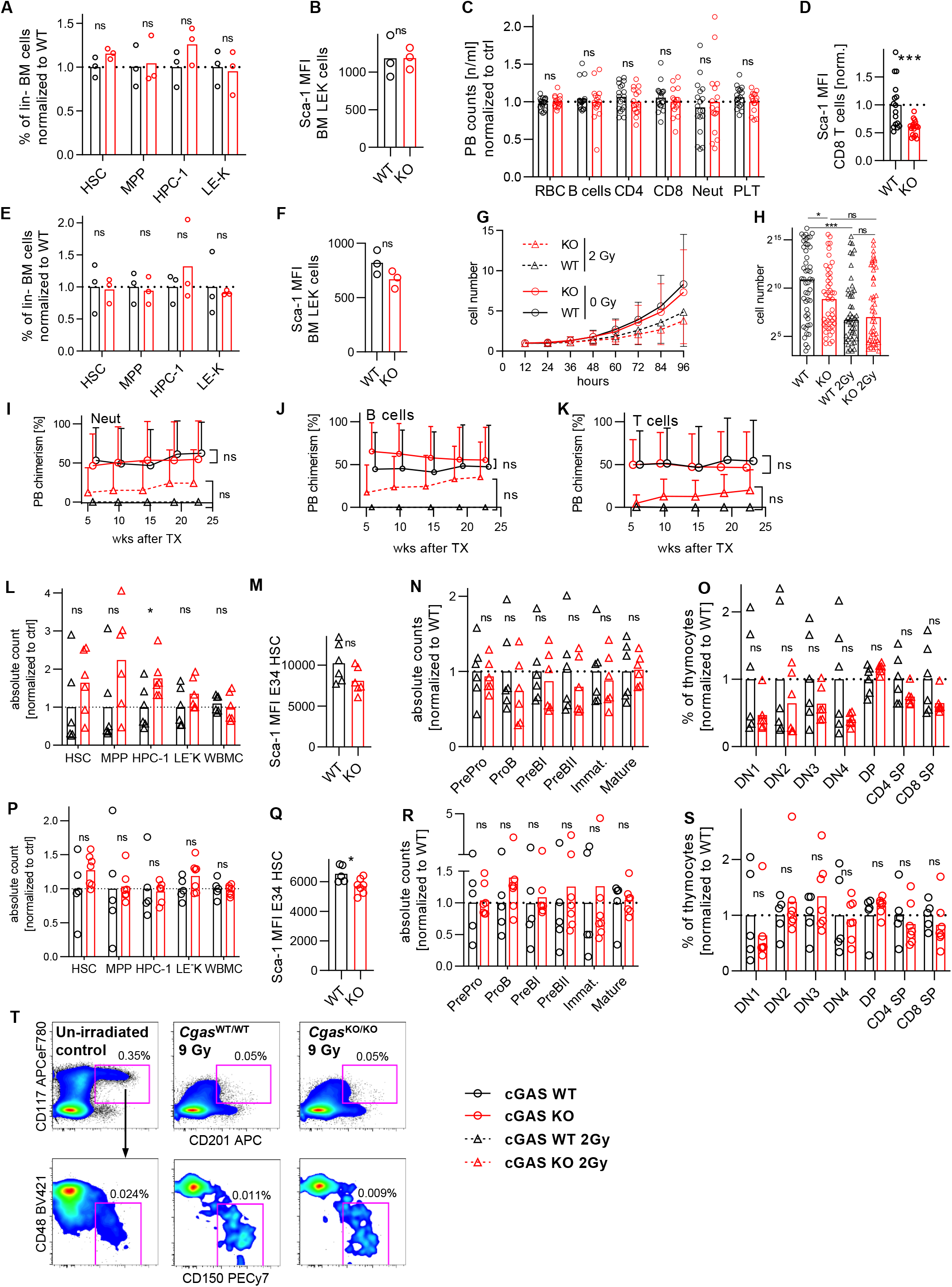
Loss of cGAS does not alter steady state or stress hematopoiesis. **A-B** BM HSPCs from *Cgas*^KO/KO^ (red) and *Cgas*^WT/WT^ (black) animals were analyzed (n=3/genotype, individual and means (bars) shown, aged 11-14 wks, significance was calculated by an unpaired Student’s t test). (A) Percentages of HSPCs among lineage negative BM cells (mean of WT control was set to 1). (B) Sca-1 expression was determined on BM LEK cells. **C-D** Peripheral blood cell counts (C) and Sca-1 expression among CD8 T cells (D) were determined (normalized to WT (set to 1), n=16-18/genotype, data from 3 independent experiments, significance was calculated by an unpaired Student’s t test, aged 9-16 wks). **E-F** (E) BM was analyzed in aged *Cgas*^KO/KO^ (red) and *Cgas*^WT/WT^ (black) animals (n=3/genotype, age 48-52 wks, significance was calculated by an unpaired Student’s t test).(E) Percentage of HSPCs among lineage negative BM (mean of WT control was set to 1). (F) Sca-1 expression was determined on BM LEK cells. **G-H** Single HSCs (LSK CD48^-/lo^CD150^+^CD34^-/lo^CD201^hi^, n=47-48/condition) isolated from aged (48-52 wks, n=3/genotype) *Cgas*^KO/KO^ (red) or WT control (black) mice were cultivated for 10 days. A fraction of cells was exposed to 2 Gy γ-radiation (triangles) before culture. (G) Cells were counted for 96h every 12h (means & SD are shown, 2-way ANOVA with Sidak post test). (H) Colony size after 10 days of culture (medians and individual colonies are shown, significance was calculated by Mann-Whitney U test). **I-K** 5 × 10^6^ whole bone marrow cells isolated from each primary recipients (Figures 6C-F) were transplanted into secondary recipients. PB neutrophil (I), B- (J) and T- (K) cell chimerism was determined (Mean and SD are shown, significance was calculated by repeated measures 2way ANOVA with Holm Sidak post test, all multiple comparisons within each time point were not significant (ns)). **L-O** *Cgas*^KO/KO^ mice (red, n=6) and *Cgas*^WT/WT^ littermate controls (black, n=6) were exposed to 2 Gy γ-radiation (same mice as Figures 6G-J). BM HSPC numbers (L), Sca-1 expression on LEK E34 HSCs (M), BM B lymphocyte progenitor numbers (N) and thymocyte frequencies (O) were determined 15 days after irradiation (individual animals and means (bars) are shown, mean of WT was set to 1 (except for M), significance was calculated by an unpaired Students t test). **P-S** *Cgas*^KO/KO^ mice (red, n=6) and *Cgas*^WT/WT^ littermate controls (black, n=6) were injected with 5-FU (same animals as in Figures 6O-R). Display of data and statistics as in Figures S8L-O. **T** *Cgas*^KO/KO^ mice and *Cgas*^WT/WT^ littermate controls were exposed to 9 Gy γ-radiation, sacrificed 10h later and BM was analyzed (same mice as in Figures 6S-T). Representative examples of gating for lineage^-^CD201^+^CD117^+^ (LEK) cells (upper row, PI^-^lineage^-^ BM cells pre-gated) and LEK CD48^-/lo^CD150^+^ cells (lower row). Percentage among total BM cells is given for each gate.

## Notes

### Competing Interest Statement

The authors have declared no competing interest.

### Summary of Updates

Main text has been revised.

