## Supplementary material for "Clearance of genome-damaged cells from the hematopoietic system via p53 without contribution by the cGAS/STING axis": Table S4 Antibodies

| **Epitope** | **Clone** | **conjugated** | **Source** |
| --- | --- | --- | --- |
| B220 | RA3-6B2 | Biotin | Biolegend |
| B220 | RA3-6B2 | FITC | Thermo Fisher |
| CD105 | MJ7/18 | APC | Thermo Fisher |
| CD117 | 2B8 | APC | Thermo Fisher |
| CD117 | 2B8 | APC-eF780 | Thermo Fisher |
| CD11b | M1/70 | APC-eF780 | Thermo Fisher |
| CD11b | M1/70 | Biotin | Thermo Fisher |
| CD150 | TC15-12F12.2 | PE-Cy7 | Biolegend |
| CD16/32 | 93 | PE-Cy7 | Thermo Fisher |
| CD16/CD32 | 93 | unconjugated | Biolegend |
| CD19 | eBio1D3 | Biotin | Thermo Fisher |
| CD19 | eBio1D3 | PE | Thermo Fisher |
| CD19 | eBio1D3 | PE-Cy7 | Thermo Fisher |
| CD201 | eBio1560 | APC | Thermo Fisher |
| CD201 | eBio1560 | PerCP-eF710 | Thermo Fisher |
| CD25 | PC61.5 | PE-Cy7 | Thermo Fisher |
| CD3 | 145-2C11 | Biotin | Thermo Fisher |
| CD3 | eBio500A2 | PE | Thermo Fisher |
| CD34 | RAM34 | eF450 | Thermo Fisher |
| CD4 | GK1.5 | Biotin | Thermo Fisher |
| CD4 | GK1.5 | PE | Thermo Fisher |
| CD41 | eBioMWReg30 | PE | Thermo Fisher |
| CD44 | IM7 | eF450 | Thermo Fisher |
| CD45.1 | A20 | APC | Biolegend |
| CD45.2 | 104 | FITC | Biolegend |
| CD48 | HM48-1 | BV421 | BD |
| CD48 | HM48-1 | PE | Thermo Fisher |
| CD49b | DX5 | Biotin | Thermo Fisher |
| CD71 | RI7217 | FITC | SouthernBiotech |
| CD8 | 53-6.7 | APC | Thermo Fisher |
| CD8 | 53-6.7 | Biotin | Thermo Fisher |
| CD8 | 53-6.7 | PE-Cy7 | Thermo Fisher |
| Gr-1 | RB6-8C5 | Biotin | Thermo Fisher |
| Gr-1 | RB6-8C5 | eF450 | Thermo Fisher |
| IgD | 11-26 | eF450 | Thermo Fisher |
| IgM | II/41 | APC-eF780 | Thermo Fisher |
| NK1.1 | PK136 | Biotin | Thermo Fisher |
| Sca-1 | D7 | FITC | Thermo Fisher |
| Sca-1 | D7 | PE | Thermo Fisher |
| Sca-1 | D7 | V500 | BD |
| Ter119 | TER-119 | Biotin | Biolegend |
| Ter-119 | TER-119 | eFluor450 | Thermo Fisher |
| Phosphor (Ser139) H2A.X | 20E3 | Unconjugated | Cell Signaling Technology |
| Goat Anti-rabbit IgG (H+L) | polyclonal | Alex Fluor 488 | Thermo Fisher |

**Table S4 Overview of employed antibodies**

Reactivity against mouse epitopes, if not otherwise mentioned.
